# ER-PM contacts regulate apical domain formation in hepatocytes

**DOI:** 10.1101/2020.04.23.057521

**Authors:** Gary Hong Chun Chung, Jemima J. Burden, Maëlle Lorvellec, Paul Gissen, Christopher J. Stefan

**Author notes:** Correspondence: Jemima J. Burden, Christopher J. Stefan. These authors contributed equally to this study (Gary Hong Chun Chung, Jemima J. Burden).

## Abstract

Apico-basal membrane polarity is fundamental for epithelial cell development and function. Polarity factors including the small GTPase Cdc42, the Par3/Par6/aPKC complex, and cytoskeletal proteins are recruited by the anionic lipids phosphatidylinositol 4,5-bisphosphate and phosphatidylserine. But how these lipids accumulate at polarised sites remains unclear. We have examined roles of contacts between the endoplasmic reticulum and plasma membrane (ER-PM contacts) in generating lipid gradients during apical domain formation. Comprehensive electron microscopy analyses in hepatocytes and epithelial spheroids revealed two distinct ER-PM contact architectures that are spatially linked to apical and baso-lateral domains. Moreover, apical domain formation was delayed in HepG2 cells upon modulating the ER-PM contact proteins E-Syt1 and ORP5. We propose ER-PM contacts regulate apico-basal polarity via the lipid transfer proteins E-Syt1 and ORP5. Importantly, our findings suggest that the spatial organisation of ER-PM contacts is a conserved feature of polarised epithelial cells.

## Introduction

The liver is composed mostly of polarised epithelial cells known as hepatocytes. Hepatocytes perform vital functions such as bile production and secretion, energy homoeostasis (glucose and lipid metabolism) and detoxification of blood (Najjar and Perdomo, 2019; Schulze, et al., 2019; Rui, 2014). These diverse biological functions involve complex membrane trafficking pathways that deliver or internalise specific cargos at designated plasma membrane (PM) domains (Schulze, et al., 2019; Gissen and Arias, 2015; Treyer and Musch, 2013). For instance, the apical domains of hepatocytes form bile canaliculi and are sites for polarised delivery of bile salt and lipid transporters, whereas the baso-lateral domains of hepatocytes are exposed to blood for insulin signalling and nutrient (glucose and lipid) uptake (Najjar and Perdomo, 2019; Schulze, et al., 2019; Gissen and Arias, 2015; Musch, 2014; Rui, 2014; Treyer and Musch, 2013). Defects in hepatocyte polarity are associated with various physiological disorders such as bile secretory failure (cholestasis), hepatic carcinoma progression, and hepatitis viral entry (Lu, et al., 2018; Key, et al., 2017; Gissen and Arias, 2015; Schulze, et al., 2012; Wang and Boyer, 2004). In addition, the liver is a highly regenerative organ involving hepatic progenitor cell proliferation and development. Therefore, it is important to understand how cell polarity and organisation are established and maintained in hepatocytes.

Several conserved proteins are essential for apico-basal polarity. Cdc42, a member of the Rho family of small GTPases, controls polarised growth in yeast and mammalian epithelial cells via the organisation of cytoskeletal and vesicular trafficking proteins (Pichaud, et al., 2019; Campanale, et al., 2017; Chiou, et al., 2017). The anterior PAR protein complex (PAR3/PAR6/aPKC) interacts with Cdc42 and additional polarity factors that regulate cell junctions and physically define the apico-basal axis in multiple cell types (Peglion and Goehring, 2019; Pichaud, et al., 2019; Kemphues, 2000). These polarity effectors are recruited and regulated, at least in part, by anionic lipids localised at the PM. For example, phosphatidylserine (PS) is required for the polarised localisation of Cdc42 (Fairn, et al., 2011). Likewise, the phosphoinositide isoform phosphatidylinositol 4,5-bisphosphate, also known as PI(4,5)P_2_, binds the polybasic C-terminus of Cdc42 (Johnson, et al., 2012; Harlan, et al., 1994). PI(4,5)P_2_ is also proposed to control the apico-basal polarity axis in epithelial cells by recruiting the PAR protein complex (Claret, et al., 2014; Shewan, et al., 2011; Pinal, et al., 2006). Accordingly, both PS and PI(4,5)P_2_ are proposed to be enriched at sites of polarised growth where they may serve as spatial landmarks for polarity factors (Hammond and Hong, 2018; Haupt and Minc, 2017; Fairn, et al., 2011; Shewan, et al., 2011; Garrenton, et al., 2010; Pinal, et al., 2006). However, while it is becoming clear that PS and PI(4,5)P_2_ serve as polarity cues, little is known regarding how membrane lipid gradients are generated and maintained at polarised sites.

Lipid transfer proteins (LTPs) that function at ER-PM contacts have been proposed to regulate PI(4,5)P_2_ and PS homeostasis. For instance, the extended synaptotagmin protein 1 (E-Syt1) has been implicated in PI(4,5)P_2_ replenishment at the PM following intense phospholipase C-mediated signalling events (Chang, et al., 2013). However, a clear role for E-Syt1 in PI(4,5)P_2_ synthesis has not been elucidated and is even controversial (Saheki, et al., 2016). In addition, the oxysterol-binding protein related protein 5 (ORP5) has been shown to deliver PS from the ER to the PM in exchange for phosphatidylinositol 4-phosphate (PI4P) at ER-PM contacts (Sohn, et al., 2016; Chung, et al., 2015). Because PI4P is the precursor for PI(4,5)P_2_ synthesis, ORP5 PS/PI4P exchange activity also modulates PI(4,5)P_2_ levels at the PM (Sohn, et al., 2018). Hence, we hypothesised that distinct LTP activities at different ER-PM contacts may fine-tune the levels and localisation of PS and PI(4,5)P_2_ at the PM. Accordingly, distinct ER-PM contacts with different physical attributes and functions may specify apical domain formation during epithelial cell development.

Here, we perform a quantitative electron microscopy (EM) analysis of hepatocytes in mouse liver and find distinct classes of ER-PM contacts. One group of ER-PM contacts have a punctate morphology and are associated with microvilli in the apical domain, whereas a second group of ER-PM contacts are extensive in size and preferentially observed at the adherens zona (lateral domains that form cell-cell contacts) neighbouring the apical domain. We observe the same patterns in mouse renal inner medullary collecting duct (mIMCD) spheroids, suggesting that ER-PM contact organisation and distribution are conserved in polarised epithelial cells. Furthermore, we demonstrate delayed apical domain formation and reduced apical domain size upon overexpressing variants of E-Syt1 and ORP5 in low-density HepG2 cultures. Our findings suggest conserved roles of ER-PM contacts in directing apical domain formation and specification in epithelial cells.

## Results

### Quantitative ultrastructure analysis of ER-PM contacts in hepatocytes

To examine the quantity, size, and distribution of ER-PM contacts in polarised epithelial cells, we conducted a comprehensive EM analysis of ER-PM contacts in hepatocytes from mouse liver (Figure 1 and Figure 1 – Figure supplement 1). The PM of the hepatocyte was classified into three domains (Figure 1a). The apical domain forms the bile canaliculi (BC) that are confined by the tight junctions; the basal domain refers to the PM exposed to the Space of Disse and the sinusoid; the lateral domain, or adherens zona, refers to the cell-cell contact zone between two hepatocytes (H/H contact). Figure 1a shows the average proportion (length %) of different PM domains in murine hepatocytes. The PM is partitioned into approximately equal portions of the basal domain (40%) and H/H contact zone (47%), whereas only 13% of the PM is apical.

**Figure 1.**
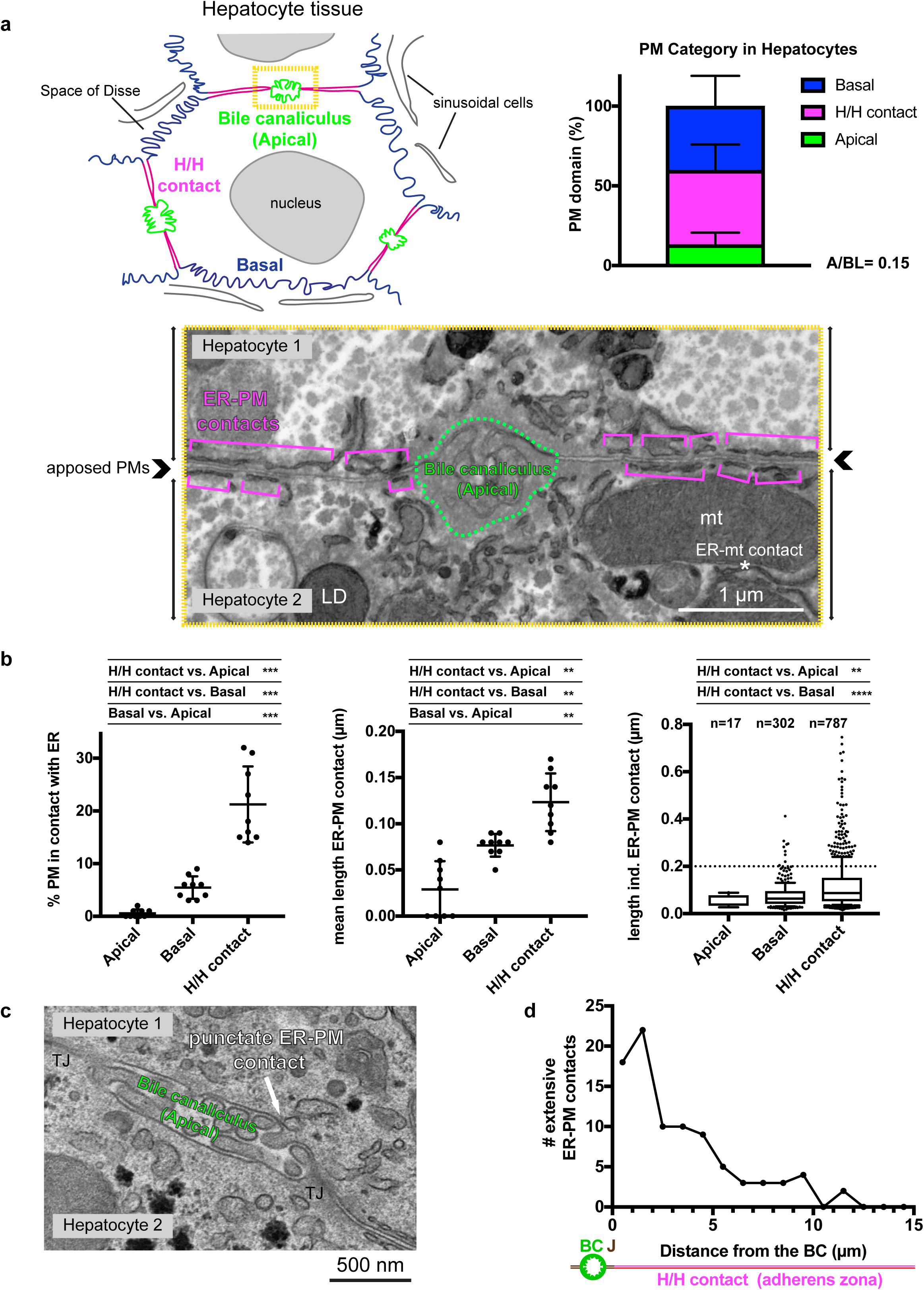
The quantity and length of ER-PM contacts in hepatocytes are PM domain-specific. **(a)** An annotated diagram of hepatocyte tissue. The apical PM domain that forms bile canaliculi (BC), adherens zona (H/H contact), and basal PM domain are colour coded in green, magenta and blue respectively. The bar chart on the right displays the proportion of each PM domain by perimeter (mean ± SD), derived from EM image analyses of 9 hepatocytes from 3 animals (see Figure 1 - Figure supplement 1 for the analysis workflow). The electron micrograph is a representative example of the apical domain and adherens zona in two apposed hepatocytes, corresponding to the yellow box of (a). The green dashed line marks the region of the BC formed by the apposing apical domains and the magenta brackets indicate the ER-PM contacts in the adherens zona (H/H contact). **mt:** mitochondria; **LD**: lipid droplet; **asterisk** (*): ER-mitochondria contact. **(b)** Quantitative analyses of ER-PM contacts in hepatocytes. The scatter dot plots show the percentage of the PM in contact with the ER (left panel) and the average length (middle panel) of ER-PM contacts in different PM domains. Data are shown as mean ± SD and were analysed by one-way ANOVA (Tukey’s multiple comparison test: ***p<0.001; **p<0.01). The box and whisker plot (right panel) depicts the median length and 10 – 90 percentiles of individual ER-PM contacts in the different PM domains of hepatocytes. We defined ER-PM contacts with length >0.2 *μ*m as “extensive ER-PM contacts”. **(c)** An example of a punctate ER-PM contact in the apical domain is shown. **(d)** The frequency distribution of extensive ER-PM contacts from the boundary of the BC (0 *μ*m; defined by the end of the junction, J) to the distal end of the adherens zona up to 15 *μ*m. Data summarise distribution of the extensive ER-PM contacts from both sides of the BC (N=22).

Next, we compared the extent of ER-PM contacts in each PM domain. The workflow for the quantitative analysis of ER-PM contacts is summarised in Figure 1 – Figure supplement 1. Approximately 20% of the PM is associated with the ER in the H/H contact zone; such coverage is 4 times and 40 times more than that found in the basal (5.4%) and the apical (0.6%) domain respectively (Figure 1b, left panel). Moreover, the average length of ER-PM contacts in the H/H contact zone is 0.12 μm, which is significantly longer than that found in the basal (0.08 μm) and the apical (0.03 μm) domains (Figure 1b middle panel). An example of a punctate ER-PM contact in the apical domain is shown in Figure 1c. Additional examples of ER-PM contacts in the apical, H/H contact, and basal domains are provided in Figure 1 – Figure supplement 2. While the apical domain displayed the lowest degree of ER-PM contact formation, the sub-apical cytoplasm is not devoid of the ER network; ER-vesicle, ER-mitochondrial, and ER-lipid droplet contacts are observed in the sub-apical cytoplasmic region (Figure 1a and Figure 1 - Figure supplement 2). The reduced number and length of ER-PM contacts at the apical PM may be due to the presence of a thick actin cortex at the apical membrane. A similar pattern was observed at the hepatocyte-cholangiocyte interface, where ER-PM contact-free zones in the basal PM correlated with the presence of a thick actin cortex (Figure 1 – Figure supplement 3). Taken together, the initial results of the EM analyses suggest the morphology of ER-PM contacts is PM domain-specific.

To gain further insight into the arrangement and features of ER-PM contacts, we plotted the length of individual ER-PM contacts in each of the PM domains. A subpopulation of extra-long ER-PM contacts greater than 0.2 μm was predominantly found in the H/H contact zone, as shown by the box and whisker plot (Figure 1b, right panel) and histogram (Figure 1 – Figure supplement 1), We have termed these extra-long ER-PM contacts as “extensive ER-PM contacts” (additional examples can be seen in Figure 1 – Figure supplement 1). Interestingly, most (∼ 60%) of the extensive ER-PM contacts locate within 2 μm of the BC, and there is a steep drop in the number of extensive ER-PM contacts as the distance from the tight junction increases (Figure 1d). These observations suggest that the formation and distribution of extensive ER-PM contacts is not a random event but under precise regulation.

### 3D reconstruction of ER-PM contacts in hepatocytes

Our 2D EM analysis indicated that distinct ER-PM contacts are formed in different PM domains. To confirm our initial observations, we examined the size and distribution of ER-PM contacts in 3D reconstructions of serial EM sections (Figure 2). This allowed a more detailed analysis of ER-PM contact morphological features such as area and shape. The PM (cyan), the ER (yellow), and ER-PM contacts (magenta) were manually traced in *xy* in each section, reconstructed into 3D models, and displayed as *xz* projections (Figure 2). 3D renderings are available in Supplemental Movies 1 and 2. Additional examples of ER-PM contact reconstructions are provided in Figure 2 – Figure supplement 1. ER-PM contacts were rare in the apical domain, but punctate ER-PM contacts were observed at the base of microvilli (Figure 2a; asterisks*). ER-PM contacts in the H/H contact domain were large and extensively occupied the associated PM (Figure 2a; chevrons >>). Some ER-PM contacts in this region even extended throughout the entire *z* series. Accordingly, the morphology of the cortical ER resembled sub-surface cisternae in the H/H contact region (Supplemental Movie 1). The basal domain had a mixed population of ER-PM contacts, as both punctate and extensive contacts were observed (Figure 2b, # marks an example). However, ER-PM contacts formed by cisternal ER in the basal domain were often smaller in size than those found in the H/H domain (Figure 2a and 2b). Interestingly, punctate ER-PM contacts were often in the vicinity of protrusions and intrusions of the basal PM (Supplemental Movie 2), similar to punctate ER-PM contacts at the base of microvilli in the apical domain.

**Figure 2.**
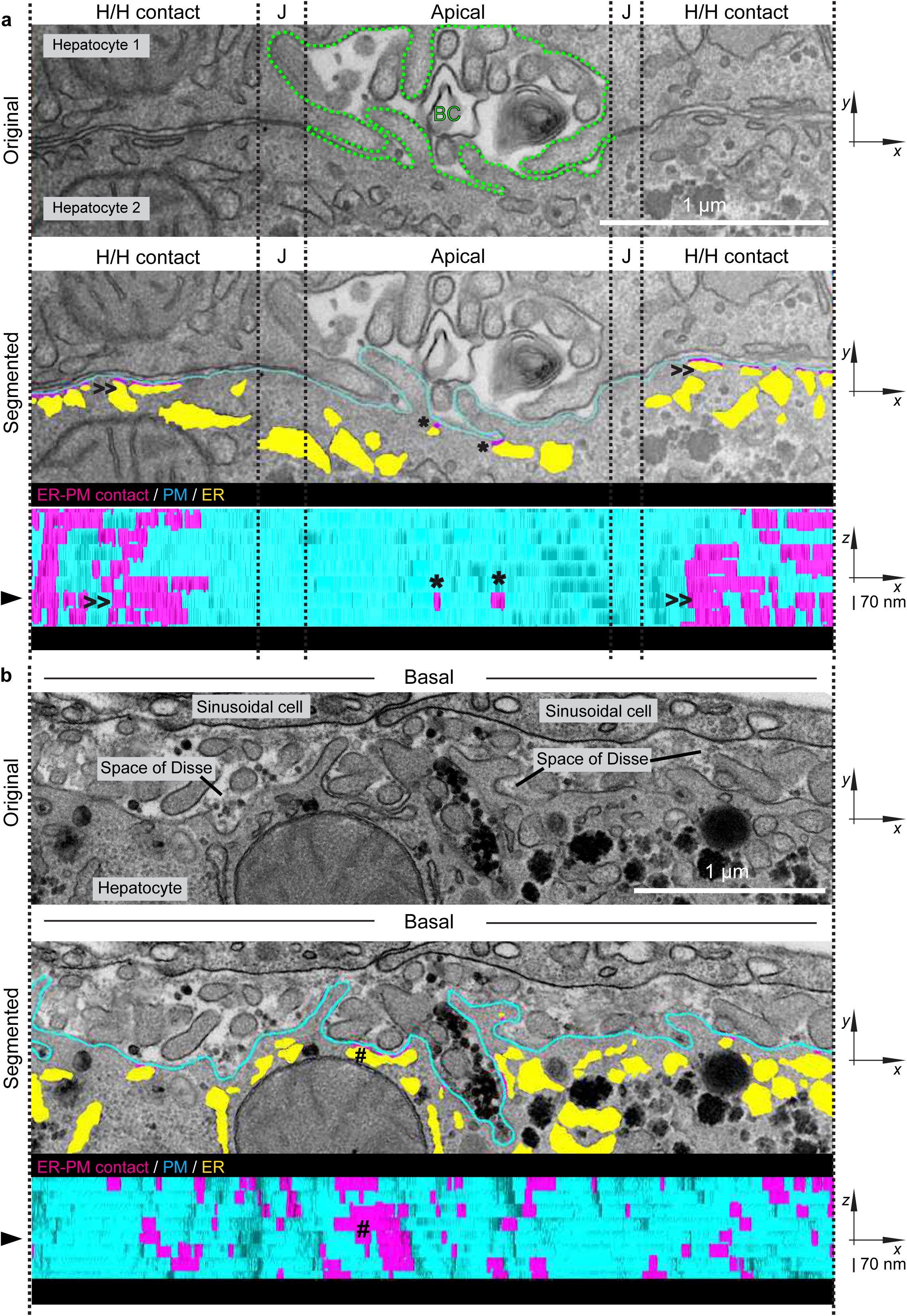
Segmentation analysis of ER-PM contacts in EM serial sections reveals their distinctive morphology in different PM domains. The PM, ER, and ER-PM contacts in **(a)** the apical and H/H contact domains and **(b)** the basal domain of hepatocytes were analysed by EM serial sectioning, tracing, and 3D reconstruction. The top panel in (a) is a labelled electron micrograph of two apposed hepatocytes consisting the apical and H/H contact domains separated by the cell junctions (J). The green dashed line marks the region of the BC. The top panel in (b) shows the basal domain facing the sinusoidal cells and the Space of Disse. The middle panels (a and b) display the traces of the PM (cyan), ER (yellow) and ER-PM contacts (magenta) on the electron micrographs shown in the top panels by manual segmentation. The segmented membranes and ER-PM contacts from 8 continuous slices (70 nm thick per slice) were reconstructed to 3D models and their ultrastructure in the different PM domains can be visualised in Supplemental Movies 1 and 2. The bottom panels (a and b) display the 3D model aligned to *x-z* axes with the intracellular side facing the observer, showing only the PM (cyan) and the tethering of the ER-PM contacts (magenta). The black arrowhead indicates the position of the segmented slice (middle panel) within the 3D model, vertical scale bar = 70 nm. Asterisk (*), chevrons (>>), and hashtag (#) annotate the examples of ER- PM contacts that possess distinctive morphology in the corresponding PM domains.

Quantitative and statistical analyses revealed a prominent divergence in the size of ER-PM contacts between the different PM domains. The mean area of ER-PM contacts was significantly larger in the H/H contact domain (10,674 nm^2^) as compared to the basal (5748 nm^2^) and apical (2534 nm^2^) domains (Figure 2 – Figure supplement 2). Accordingly, the maximal ER-PM contact area in the apical domain was an order of magnitude less than the max contact area in the basal and H/H domains (see the table in in Figure 2 – Figure supplement 2). The extensive ER-PM contacts, especially those larger than 20,000 nm^2^, were predominantly found in the H/H contact domain (shown in the histogram in Figure 2 – Figure supplement 2). Overall, our quantitative 2D and 3D EM analyses demonstrate that ER-PM contacts with distinct sizes and architectures are formed and distributed in specific PM domains (Figures 1 and 2 and their Figure supplements). This suggests that distinct ER-PM tethering proteins may form morphologically distinct ER-PM contacts and may even be involved in the establishment and maintenance of apical-basolateral polarity.

### PI(4,5)P_2_ and PS are present in the apical domains of HepG2 couplets

Certain phospholipids are proposed to serve as spatial cues for apico-basolateral polarity in epithelial cells. For example, phosphoinositide lipid isoforms display asymmetrical partitioning between apical and basolateral domains to specify PM domain identity and function (Roman-Fernandez, et al., 2018; Shewan, et al., 2011; Martin-Belmonte, et al., 2007). In particular, PI(4,5)P_2_ is proposed to be enriched in the apical domain of polarised MDCK cells (Roman-Fernandez, et al., 2018; Martin-Belmonte, et al., 2007) where it serves as a key determinant in apical domain formation and function (Shewan, et al., 2011). In addition, phosphatidylserine (PS) is enriched in the cytoplasmic leaflet of the PM at sites of polarised growth in yeast and dynamic PM structures in mammalian cells (Kay and Fairn, 2019; Kay, et al., 2012; Fairn, et al., 2011). However, it is not known whether PI(4,5)P_2_ or PS are present in the apical domain of hepatocytes or whether they are involved in BC formation. Moreover, it is not known whether or how ER-PM contacts might influence apical domain and BC formation during hepatocyte development.

We used HepG2 cells as a model to monitor the localisation of PI(4,5)P_2_ and PS during hepatocyte cell polarisation and BC development. HepG2 cells have been shown to polarise and form BC-like structures between the opposing apical membranes of cell couplets (also referred to as doublets) (Gissen and Arias, 2015; Chiu, et al., 1990). We categorised BC development into three consecutive stages based on the morphology of HepG2 couplets (Figure 3a). The apposition stage refers to initial physical contact between two cells. The developing BC stage appears as a globular accumulation of apical membrane reporters at sites of PM apposition, as assessed by confocal microscopy. This may be due to crowding of microvilli between the developing apical domains of the opposing cells (see Supplemental Movie 3 for EM serial sections of a developing BC in a HepG2 couplet). A mature BC is defined by a dilated lumen and the presence of clearly resolved microvilli along the edges of hemi-canaliculi.

**Figure 3.**
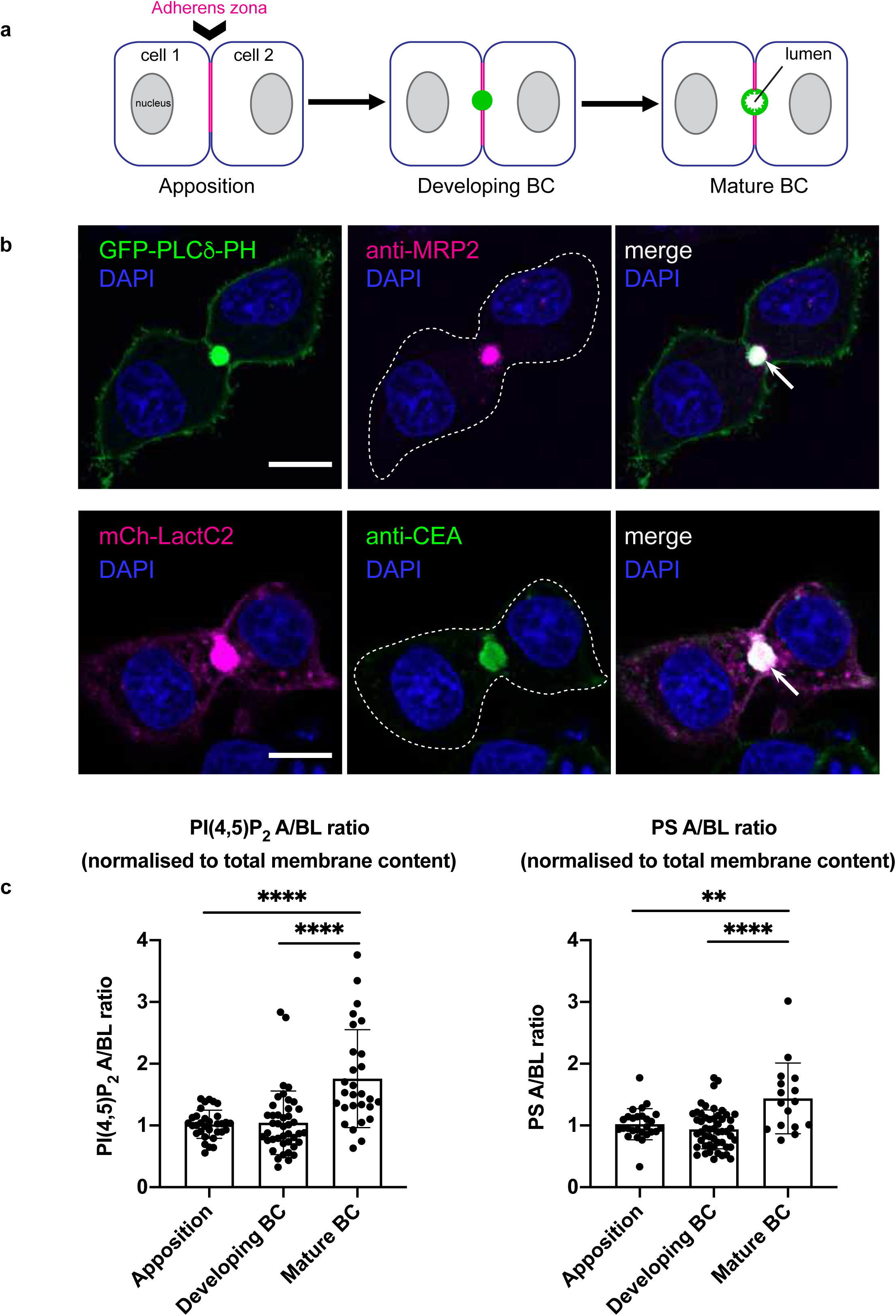
The bile canaliculi are enriched in polarised lipids PI(4,5)P_2_ and PS. **(a)** A cartoon illustrates the BC development in hepatocyte cell lines and the morphological characters associated with each stage. The basal PM, apposing PM (cell-cell contact), and apical PM (BC) are coloured in blue, magenta and green respectively. **(b)** Confocal images of HepG2 couplets in the developing BC stage. HepG2 cells overexpressing the PI(4,5)P_2_ (GFP- PLC^*δ*^-PH; green) or PS (mCherry-Lact-C2; magenta) reporter were labelled with an established BC marker (anti-MRP2 or anti-CEA antibody) made visible by applying a fluorescent secondary antibody. Nuclei were stained with DAPI (blue) and the white arrow annotates the position of the developing BC within the cell couplet. Scale bar = 10 m. **(c)** Spatial ratiometric fluorescence analysis of PI(4,5)P_2_ and PS reporters in HepG2 couplets. HepG2 cells overexpressing the lipid reporters were stained with CellMask DeepRed (a general PM dye), and the fluorescence of the lipid reporters and the dye at different PM domains was measured as illustrated in Figure 3 – Figure supplement 1b. Scatter dot plots display the apical (A) to basolateral (BL) lipid intensity (mean ± SD) throughout each BC developmental stage. The intensity of the lipid reporters has been normalised to a general membrane reporter (CellMask) in each PM domain (see Figure 3 –Figure supplement 1c). Data were analysed by one-way ANOVA followed by Tukey’s multiple comparison test: ****p<0.0001; **p<0.01. N=2 independent experiments.

Next, we expressed the biosensors GFP-PLC_*δ*_-PH and GFP-Lact-C2 in HepG2 cells to monitor PI(4,5)P_2_ and PS localisation during BC development. In control cells, the PI(4,5)P_2_ and PS reporters were observed at developing BC that co-stained with known BC marker proteins (MRP2 or CEA; Figure 3b). The PS reporter was also found at developing and mature BC that co-stained with the known apical domain protein Par3 (Figure 3 – Figure supplement 1a). We also employed a quantitative approach to assess the relative levels of lipid reporters in the apical and basolateral domains of HepG2 couplets at different developmental stages (Figure 3 – Figure supplement 1b). Ratiometric imaging using a general PM dye indicated that the PI(4,5)P_2_ and PS reporters were slightly enriched (>2-fold) in mature BC (Figure 3c and Figure 3 – Figure supplement 1c), suggesting that PI(4,5)P_2_ and PS may accumulate at the apical domain during the course of BC development. Thus, PI(4,5)P_2_ and PS may be important for apical domain formation and BC maturation in HepG2 couplets.

### Overexpression of mutant E-Syt1 proteins impairs BC formation

Next, we investigated whether proteins known to form and function at ER-PM contacts are involved in BC formation in HepG2 couplets. The extended synaptotagmin proteins (E- Syt1/2/3) have been shown to localise to ER-PM contacts where they are implicated in Ca^2+^- mediated membrane lipid dynamics (Bian, et al., 2018; Saheki, et al., 2016; Chang, et al., 2013; Giordano, et al., 2013). Their role in PI(4,5)P_2_ metabolism has been somewhat controversial. One study has implicated E-Syt1 in PI(4,5)P_2_ synthesis (Chang, et al., 2013), another has suggested a role for E-Syt2 in PI(4,5)P_2_ turnover (Dickson, et al., 2016), while yet another study found that loss of the E-Syt1/2/3 proteins had no significant effect on PI(4,5)P_2_ metabolism (Saheki, et al., 2016). We overexpressed mCherry-tagged wild type E- Syt1 or mutant forms of the protein (E-Syt1-D406A or E-Syt1-D724A bearing a substitution in C2A or C2C respectively (Chang, et al., 2013) in HepG2 cells (Figure 4a and Figure 4 – Figure supplement 1a). The SMP domain in the E-Syt proteins form dimers that enable lipid transfer between membrane bilayers (Saheki, et al., 2016; Yu, et al., 2016; Schauder, et al., 2014). Since E-Syt proteins function as obligate homo- or heterodimers, we reasoned that mutant forms defective in Ca^2+^-mediated ER-PM tethering may confer dominant negative effects when overexpressed. Expression of wild type E-Syt1 had no major effects on PI(4,5)P_2_ and PS localisation (Figure 4a) or BC development (Figure 4b) in HepG2 cells. At 24h post-transfection, approximately 65% of E-Syt1-expressing couplets were in the cell-cell apposition stage, 30% displayed developing BC, and 5% had formed mature BC (Figure 4b), similar to control cells. In HepG2 couplets expressing E-Syt1-D406A or E-Syt1-D724A, there was an increase in the apposition stage and corresponding decreases in the developing (2-fold) and mature (>3-fold) BC stages (Figure 4b, as assessed by both lipid biosensors). We also quantified the aspect ratio between apical and basolateral domains as an additional parameter to monitor the progression of BC (developmental stage and size) and cell polarity. Hepatocytes in mouse liver and mature HepG2 couplets have a similar apical to basolateral aspect ratio (A/BL ratio = 0.15; see Figure 1a and Figure 4 – Figure supplement 1b). The apical to basolateral aspect ratio was reduced in HepG2 couplets expressing the E-Syt1- D406A and E-Syt1-D724A mutant proteins, but not wild type E-Syt1, as compared to control cells (Figure 4 – Figure supplement 1b) further suggesting that BC development is delayed upon E-Syt1 impairment. Interestingly, intracellular PS-positive clusters were readily observed (often in perinuclear regions) in cells expressing the mutant E-Syt1 proteins (Figure 4a, yellow arrowheads). Taken together, these results suggest that E-Syt1 may be involved in the accumulation of PS and PI(4,5)P_2_ at the apical PM, possibly by PS transfer at ER-PM contacts (Figure 4 – Figure supplement 1c), and that E-Syt1 may be important in BC formation and development. Previous studies have shown that PS stimulates PI4P 5-kinase (PIP5K) activity and PI(4,5)P_2_ synthesis (Nishimura, et al., 2019; Fairn, et al., 2009). A role of E-Syt1 in PS regulation may explain the previously described role for E-Syt1 in PI(4,5)P_2_ synthesis (Chang, et al., 2013) (see Figure 4 – Figure supplement 1c and d). In addition, E- Syt1 may serve as a bridge or scaffold for additional proteins involved in PI(4,5)P_2_ synthesis (Chang, et al., 2013) that in turn may direct or reinforce polarised secretion of PS-containing vesicles (Fairn, et al., 2009).

**Figure 4.**
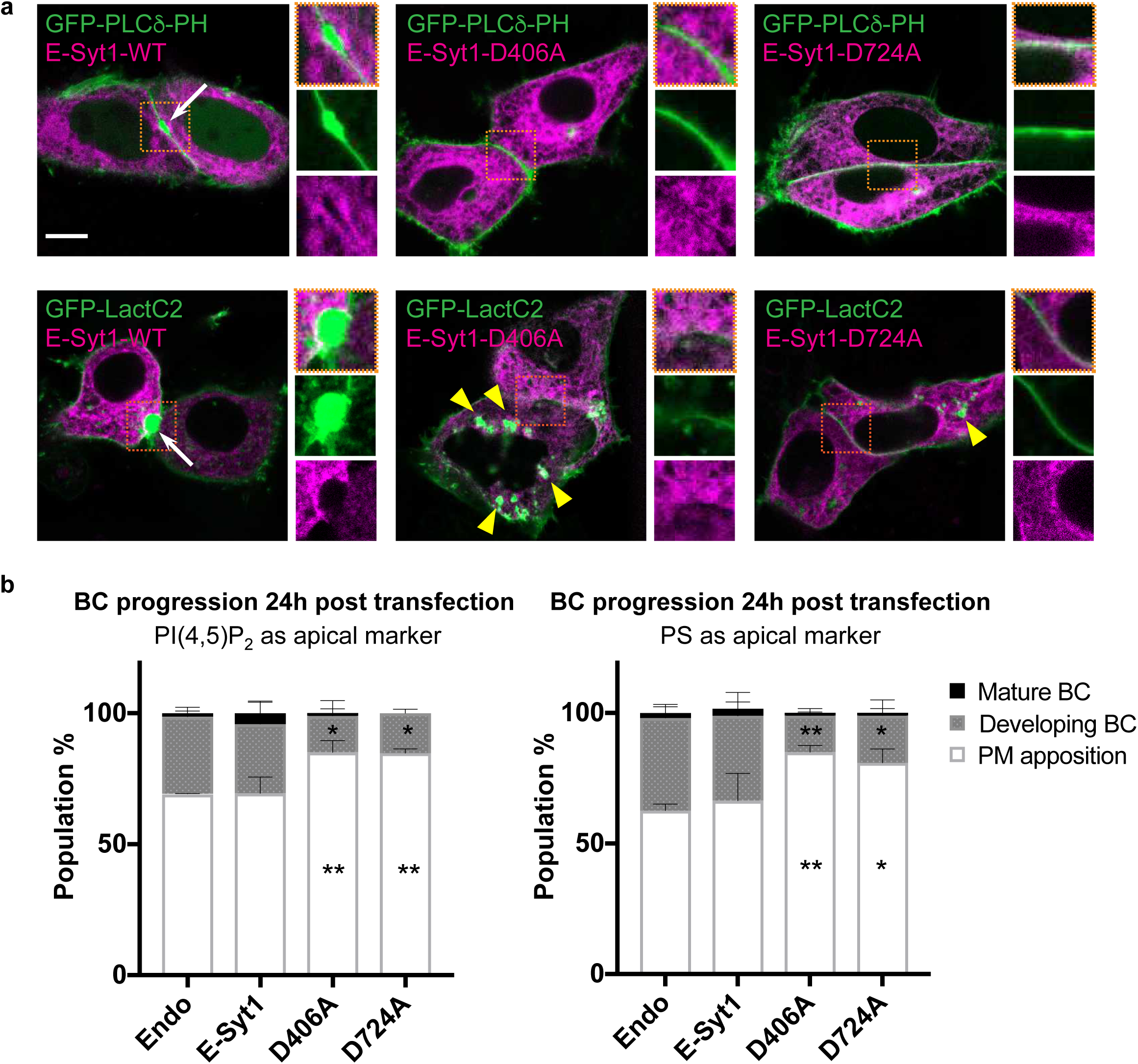
BC development is reduced in cells overexpressing E-Syt1 mutants. **(a)** Confocal microscopy images of live HepG2 couplets co-transfected with either a PI(4,5)P_2_ or PS reporter (GFP-PLC^*δ*^-PH or GFP-Lact-C2; green) and introduced mCherry-tagged E-Syt1 variants (magenta). Small panels are 2X magnified images corresponding to the region of apposition (orange box) in each cell couplet. White arrows indicate the compartment of the developing BC and yellow arrowheads indicate the PS-positive cytoplasmic puncta. Scale bar = 10 *μ*m. **(b)** Number of the transfected HepG2 couplets at each BC developmental stage, as in percentage (mean ± SD). Data from N=3 independent experiments were analysed by one-way ANOVA coupled with Dunnett’s test: **p<0.01; *p<0.05; n ≈ 80 cell couplets per condition; control cells (Endo).

### ORP5 modulates BC formation

ORP5 (also named OSBPL5) functions in PS transport at ER-PM contacts by exchanging phosphatidylinositol 4-phosphate (PI4P) for PS at the PM (Sohn, et al., 2016; Chung, et al., 2015) (see Figure 5 – Figure supplement 1). Accordingly, ORP5 has been implicated in the regulation of PS, PI4P, and PI(4,5)P_2_ levels at the PM (Sohn, et al., 2018; Ghai, et al., 2017; Sohn, et al., 2016; Chung, et al., 2015). Loss of ORP5 increases PI4P and PI(4,5)P_2_ whereas ORP5 overexpression decreases levels of these lipids at the PM. We overexpressed mCherry-tagged wild type ORP5 or a mutant form lacking the PH domain, ORP5ΔPH that does not localize to ER-PM contacts (Ghai, et al., 2017) and assessed their impact on BC development in HepG2 cells (Figure 5 and Figure 5 – Figure supplement 1a). Expression of wild type ORP5 impaired BC development as shown by both the PI(4,5)P_2_ and PS probes (Figure 5b). The fraction of cell couplets in the apposition stage increased (1.3-fold) while the fraction of the developing and mature BC stages decreased (2-fold and approximately 5-fold, respectively; Figure 5b). In contrast, expression of the mutant ORP5ΔPH protein had no major effects on PI(4,5)P_2_ and PS distribution (Figure 5a) or BC development (Figure 5b). Likewise, the apical aspect ratio was reduced in HepG2 couplets expressing the wild type ORP5 protein, but not the mutant ORP5ΔPH protein, confirming that BC formation is delayed upon ORP5 overexpression (Figure 5 – Figure supplement 1c). Thus, ORP5 activity must be precisely regulated for the proper levels and distribution of PI(4,5)P_2_ and PS during BC development.

**Figure 5.**
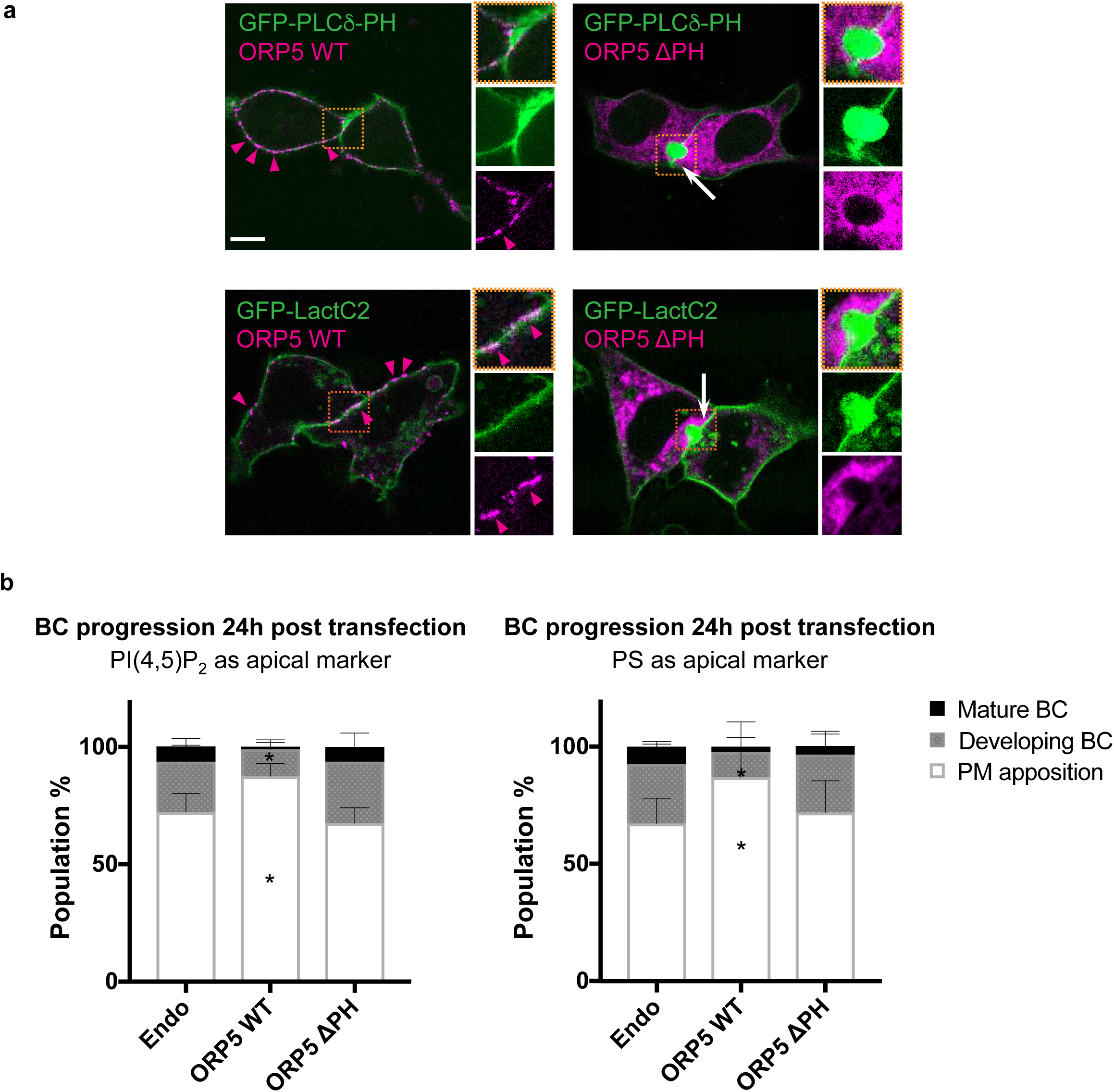
Overexpression of ORP5 reduces BC development. **(a)** Confocal microscopy images of live HepG2 couplets co-transfected with either a PI(4,5)P_2_ or PS reporter (green) and mCherry-tagged ORP5 variants (magenta). Small panels are 2X magnification of the apposed region (orange box) in each cell couplet. Magenta arrowheads indicate the localisation of WT ORP5 at the PM. White arrows indicate the position of the developing BC. Scale bar = 10 *μ*m. **(b)** Number of the transfected HepG2 couplets at each BC developmental stage (mean % ± SD), summarising observations from N=4 independent experiments for the PI(4,5)P_2_ reporter and N=3 independent experiments for the PS reporter. Data were analysed by one-way ANOVA followed by Dunnett’s test: **p<0.01; *p<0.05; control column: Endo; n ≈ 60 cell couplets per condition.

### The organisation and architecture of ER-PM contacts are conserved in epithelial organoids

To determine whether the organisation and morphology of ER-PM contacts are conserved, we examined an epithelial organoid model. We performed a quantitative EM analysis, identical to that done on the hepatocytes, on three spheroids developed from mouse renal inner medullary collecting duct (mIMCD) cells. The PM of individual cells in the spheroids was classified into three domains: the apical PM facing the enclosed lumen, the basal PM facing the external gel matrix, and the lateral PM domain that forms cell-cell contacts (C/C contact zone) (Figure 6a). The proportion of each PM domain in the IMCD spheroids was similar to hepatocytes, with an equal portion of basal (38%) and C/C domain (39%) and the apical domain in least abundance (23%). Likewise, the distribution and morphology of ER-PM contacts in mIMCD spheroids resembled those observed in hepatocytes from mouse liver. The C/C contact domain was significantly associated with the ER (8.3%, Figure 6b and see Figure 6 - Figure supplement 1a for additional examples) similar to the H/H domain in hepatocytes, and this coverage (% of PM associated with ER) was 4 times more abundant than that observed in the apical (1.7%) and the basal (1.0%) domains. The apical and basal domains showed no appreciable difference in ER-PM contact formation. This may be because the basal domains in the spheroids face the cell-free gel matrix whereas the basal domain in hepatocytes is exposed to sinusoidal cells and blood that may provide some physiological cues. ER-PM contacts in the C/C contact domain were also significantly longer than that in the apical and the basal domains (Figure 6b). In addition, a subpopulation of ER-PM contacts that meet the criteria of extensive ER-PM contacts (>0.2 μm) were observed in the C/C contact domain, as shown in the box and whisker plot in Figure 6b and the corresponding histogram in Figure 6 - Figure supplement 1. Finally, we studied the distribution of extensive ER-PM contacts. More than 80% localised within 2 μm of the apical domain just beneath the tight junction (Figure 6c). Overall, we conclude that the organisation and morphological features of ER-PM contacts, particularly the presence of extensive ER-PM contacts in the adherens zona, are conserved in epithelial cells.

**Figure 6.**
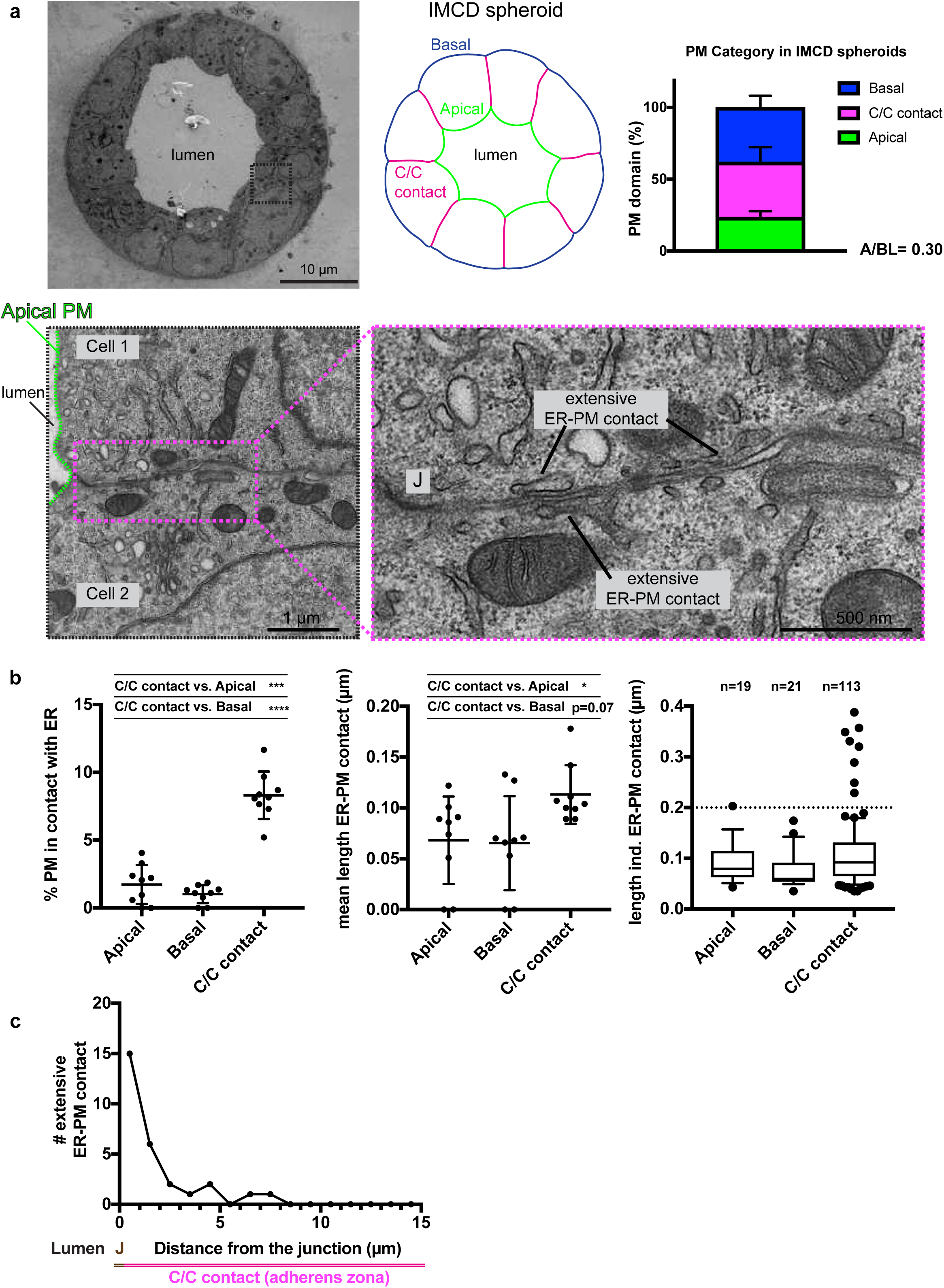
The PM domain-specific enrichment of ER-PM contacts is conserved in polarised epithelial organoid. **(a)** An electron micrograph showing a cross-section of an entire IMCD spheroid and a colour-coded diagram depicting different PM domains within the spheroid (apical = green; basal = blue; cell-cell contact/lateral = magenta). The bar chart displays the proportion of each PM domain by perimeter (mean ± SD) in polarised IMCD cells, summarising EM analyses of 9 cells from 3 spheroids. The magnified electron micrographs highlight the cell- cell contact region within the spheroid (black dashed box); apical PM is outlined in green and the extensive ER-PM contacts (length > 0.2 *μ*m) are annotated in the highest magnification. **J**: cell junction. **(b)** Quantitative analyses of ER-PM contacts in IMCD spheroids, identical to the analyses described in Figure 1b. Data of the scatter dot plots were analysed by one-way ANOVA followed by a Tukey’s multiple comparison test: ****p<0.0001; ***p<0.001; *p<0.1. **(c)** The frequency distribution of extensive ER-PM contacts from the boundary of the lumen (0 *μ*m = cell junction) to the distal end of the adherens zona up to 15 *μ*m, data gathered from 4 spheroids consisting 41 cells.

## Discussion

### The organisation of ER-PM contacts is conserved in polarised epithelial cells

Despite the discovery of ER-PM contacts decades ago (Henkart, et al., 1976; Rosenbluth, 1962; Porter and Palade, 1957) there is no thorough characterisation of ER-PM contacts in polarised epithelial cells. While the anatomical detail of junctional complexes in epithelial cells (Farquhar and Palade, 1963) and bile canaliculi in hepatocytes (Jeejeebhoy, et al., 1980; Phillips, et al., 1976; Oda, et al., 1974; Phillips, et al., 1974) have been described, the morphology of ER-PM contacts has not been quantitatively examined. Our EM analysis provides detailed information on the organisation of ER-PM contacts in distinct PM domains (apical and basolateral) of hepatocytes, HepG2 couplets, and mIMCD spheroids. We found striking differences in the quantity, size, and architecture of ER-PM contacts between the different PM domains. ER-PM contacts are scarce at the apical PM but are highly abundant at the lateral domains where cell-cell junctions form (Figures 1 and 6). Previous EM studies have mentioned an “organelle-free” sub-canalicular zone (Jeejeebhoy, et al., 1980; Phillips, et al., 1976; Oda, et al., 1974; Phillips, et al., 1974). The scarcity of ER-PM contacts at the apical PM may be due to the presence of a thick actin cortex (Figures 1 and 2). Accordingly, recent studies have reported coordination between ER-PM contacts and the actin cytoskeleton (Hsieh, et al., 2017; van Vliet, et al., 2017; Hartzell, et al., 2016). In contrast, there is an extensive system of ER-PM contacts at the lateral domains of hepatocytes and mIMCD spheroids. This pattern in ER-PM contact distribution may even be conserved throughout evolution. ER-PM contact anisotropy has been described in budding and fission yeast where sites of polarised growth have fewer ER-PM contacts compared to mother cells or the lateral PM domains respectively, particularly in G1 phase of the cell cycle when polarised growth is most prominent (Ng, et al., 2018; West, et al., 2011).

### Distinct ER-PM contacts are observed in polarised epithelial cells

ER-PM contacts in the apical and the lateral domains (adherens zona) of epithelial cells display distinct architectures. At the apical domain, punctate ER-PM contacts are formed by cortical ER tubules and extensive ER-PM contacts are formed in the lateral domain by flat subsurface ER cisternae (Figure 2 and Supplemental Movie 1). The punctate ER-PM contacts are often found at the base of microvilli in the apical domain or occasionally at the base of highly curved membranes in the basolateral domains, suggesting a relationship with local PM curvature. Similarly, punctate ER-PM contacts have been observed at the base of microvilli in *Drosophila* photoreceptor cells (Suzuki and Hirosawa, 1994). In contrast, the extensive ER-PM contacts are associated with planar lateral PM domains. The majority of the extensive ER-PM contacts localise just outside the boundary of the apical domains (within 2 microns of the tight junctions; Figure 1c and Figure 6c), implying they may function to define the apical boundary and tight junction positioning or they may regulate junctional complexes in the lateral domains.

Specialised ER-PM contacts with distinct features may not be limited to epithelial cells. In neurons, punctate ER-PM contacts are present at the pre-synaptic terminal (Hayashi, et al., 2008) whereas ER-PM contacts appear cisternal in other regions such as the soma and the base of axons and dendrites (Sun, et al., 2019; Rosenbluth, 1962). The existence of both punctate and extensive ER-PM contacts suggests structural plasticity and specialisation at designated PM domains. Accordingly, distinct proteins or complexes may form the punctate and the extensive ER-PM contacts. Consistent with this idea, the tricalbin proteins (E-Syt orthologs in yeast cells) form regions of extreme curvature at the cortical ER (Collado, et al., 2019; Hoffmann, et al., 2019), suggesting that E-Syt proteins may be present at punctate ER-PM contacts. In contrast, the ER-localised VAP (VAMP-associated proteins) orthologs Scs2/22 are associated with more extensive ER-PM contacts formed by cisternal ER in yeast (Collado, et al., 2019; Hoffmann, et al., 2019). Likewise, distinct ER-PM structures have been observed by cryo-electron tomography in primary neurons (Fernandez-Busnadiego, et al., 2015).

### E-Syt1 promotes apical domain formation

PI(4,5)P_2_ has been proposed to control apical domain formation and identity in polarised epithelial cells (Shewan, et al., 2011). We used HepG2 cells as a model to address whether ER-PM contacts are involved in PI(4,5)P_2_ regulation at the apical domain. We confirmed the presence of PI(4,5)P_2_ as well as PS in mature bile canalicular (BC) membranes of HepG2 couplets using lipid biosensors and ratiometric analyses. Previous work has demonstrated that PS can activate PI4P 5-kinase (PIP5K) for PI(4,5)P_2_ synthesis (Nishimura, et al., 2019; Fairn, et al., 2009), suggesting that these two lipids are spatially and functionally related. Our findings indicate a role for E-Syt1 in the regulation of PS and PI(4,5)P_2_ during apical domain formation and maturation (Figure 4). But how might E-Syt1 regulate PS distribution and what are the roles for PS in apical domain formation and BC maturation? Our results suggest that E-Syt1 may deliver PS from the ER to the PM, as the PS reporter accumulated on intracellular clusters in cells expressing the mutant proteins. The SMP domain of the E-Syt proteins form dimers and transfer glycerolipids including PS *in vitro* (Bian, et al., 2018; Saheki, et al., 2016; Yu, et al., 2016; Schauder, et al., 2014). Thus, the E-Syt proteins may transfer PS directly from the ER to the PM. However, both vesicular and non-vesicular transport mechanisms are implicated in the control of PS at the PM in yeast cells (Nishimura, et al., 2019; Moser von Filseck, et al., 2015; Fairn, et al., 2011). Accordingly, the E-Syt proteins may transfer PS from the ER to secretory vesicles that subsequently traffic to the apical domain. Consistent with this, ER-vesicle contacts are observed in the sub-apical cytoplasmic region (Figure 1a and Figure 1 – Figure supplement 2). Alternative roles for E- Syt1 in PS regulation may exist. For example, the E-Syt proteins have been implicated in PS asymmetry in the PM bilayer (Bian, et al., 2018). PS is normally enriched in the cytoplasmic leaflet of the PM, but PS levels increase in the extracellular leaflet upon loss of E-Syt1 function (Bian, et al., 2018). In addition, E-Syt1 is involved in recycling diacylglycerol from the PM to the ER (Saheki, et al., 2016). Diacylglycerol is used for phosphatidylcholine (PC) synthesis in the ER via the Kennedy pathway, and PC is used as a substrate for generation of PS in the ER by PS synthase 1 (Nishimura and Stefan, 2020). Possibly, diacylglycerol pools may become limiting in the ER upon impaired E-Syt1 function and this could result in reduced PS synthesis. However, the PS reporter did not appear to display increased cytoplasmic localisation in cells expressing the mutant E-Syt1 proteins. Instead, the PS reporter accumulated on intracellular compartments.

Tethering of E-Syt1 to the PM is PI(4,5)P_2_- and Ca^2+^-dependent via its C2 domains, (Idevall-Hagren, et al., 2015; Giordano, et al., 2013). Interestingly, upon addition of Ca^2+^ to the extracellular medium, isolated MDCK cells form cell-cell junctions where local intracellular Ca^2+^ levels are dramatically increased (Nigam, et al., 1992). Possibly, E-Syt-mediated ER-PM contacts may form in response to local Ca^2+^ signals at initial sites of cell-cell apposition. In line with this notion, we observed ‘hotspots’ of PS at sites of cell-cell apposition in HepG2 couplets (Figure 4 – Figure supplement 1a). Accordingly, E-Syt-mediated ER-PM contacts could result in localised PS accumulations that promote PIP5K activity and PI(4,5)P_2_ synthesis that in turn trigger more E-Syt1-mediated ER-PM contacts during early stages of cell polarisation (Figure 7a). Together, PS and PI(4,5)P_2_ may then subsequently recruit polarity effectors such as Cdc42 and the PAR protein complex to direct actin cytoskeletal dynamics that coalesce individual mesoscale membrane environments into a single apical domain (Figure 7a). Once apical identity is established, polarised secretion and actin polymerisation may drive microvilli formation during apical domain development. Upon apical domain maturation and formation of a thick sub-apical actin cortex, ER-PM contacts appear to be largely excluded from the apical domain, but E-Syt-mediated ER-secretory vesicle contacts and polarised secretion may then contribute to the maintenance of PS and PI(4,5)P_2_ levels at the mature apical domain (Figures 7a and 7b). Actin cytoskeletal dynamics and polarised secretory events taking place in the maturing apical domain may also coincide with the formation of extensive ER-PM contacts in the lateral domains (Figure 7b).

**Figure 7.**
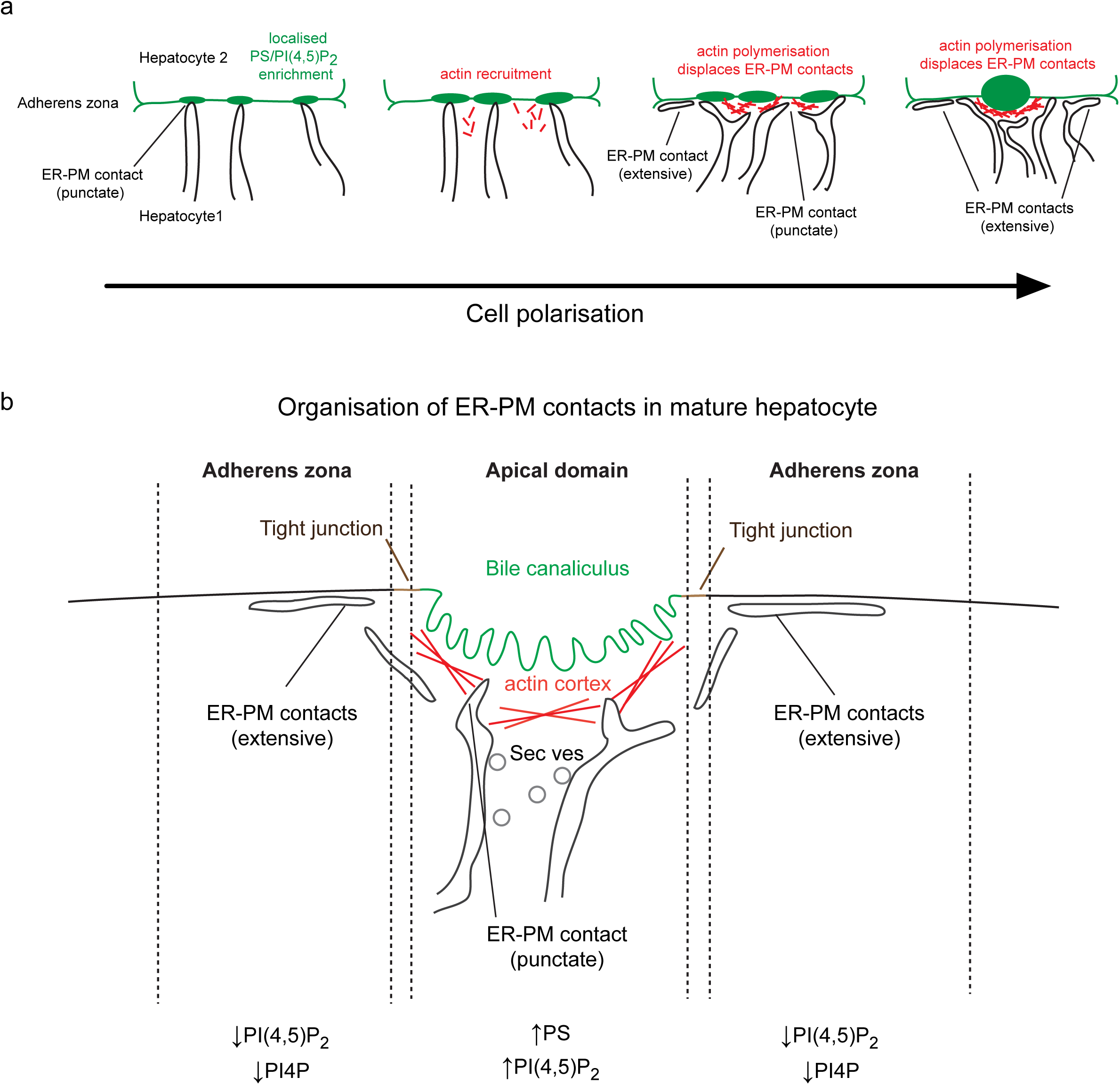
A speculative model for distinct ER-PM contacts during apical domain formation. **(a)** E-Syt-mediated ER-PM contacts promote local enrichment of PI(4,5)P_2_ and PS at the early stage of cell polarisation; **(b)** ORP-mediated ER-PM contacts maintain the lipid gradient at the polarised site by limiting the PI4P/PI(4,5)P_2_ in the adherens zona, possibly setting up a boundary for the apical domain during polarisation and maturation.

It should be noted that our results did not demonstrate a requirement for E-Syt1 in BC formation and maturation *per se*. Rather, BC development was delayed in HepG2 cells expressing mutant E-Syt1 proteins. Intriguingly, loss of all three E-Syt proteins has no apparent effect in the mouse (Sclip, et al., 2016; Tremblay and Moss, 2016). However, up-regulation of other lipid transfer proteins or polarised secretion may compensate for the loss of the E-Syt proteins. Since the E-Syts form Ca^2+^-inducible ER-PM contacts, it may be interesting to examine E-Syt knock out hepatocytes upon membrane stress induced by a high fat or glucose diet or conditions that mimic cholestasis.

### ORP5-mediated ER-PM contacts define apical domain size

While punctate E-Syt-mediated ER-PM contacts and polarised secretion may establish lipid gradients in the apical domain, additional mechanisms must be in place to restrict or retain the lipid gradients upon apical domain maturation. Membrane curvature and protein assemblies may reduce lipid diffusion in the microvilli-lined apical domain. Repulsive forces between anionic lipids including PS are proposed to induce membrane curvature (Hirama, et al., 2017), and membrane curvature can in turn decrease lipid diffusion rates (Hansen, et al., 2019; Trimble and Grinstein, 2015). Moreover, PS and PI(4,5)P_2_ recruit and regulate Rho family GTPases and cytoskeletal proteins that form actin- and septin-based assemblies that in turn may confine PS and PI(4,5)P_2_ in the apical domain (Kay and Fairn, 2019; Trimble and Grinstein, 2015; Kay, et al., 2012; McLaughlin and Murray, 2005). In addition, tight junctions (TJs) form paracellular diffusion barriers between polarised epithelial cells. However, although TJs can restrict the diffusion of lipids in the extracellular leaflet of the PM, this barrier does not extend to the inner leaflet (Trimble and Grinstein, 2015; Dragsten, et al., 1982). Thus, anionic lipids that diffuse into (or are delivered to) the apical domain may be confined in this region by geometric and electrostatic mechanisms to promote apical domain formation. Yet little is known about regulatory mechanisms that restrict apical domain size or define its boundaries set by TJ positioning.

While E-Syt1 appears to facilitate apical domain formation, ORP5 activity instead opposes or restricts apical domain development (Figure 5) and size (apical aspect ratio, Figure 5 – Figure supplement 1). Accordingly, ORP5 overexpression impaired PS and PI(4,5)P_2_ accumulation at the apical domain. ORP5 carries out PI4P/PS exchange at ER-PM contacts, and overexpression of ORP5 results in lower PM PI4P whereas ORP5 knockdown increases PM PI4P (Sohn, et al., 2016; Chung, et al., 2015). Extraction of PI4P from the PM by ORP5 may attenuate PIP5K activity and PI(4,5)P_2_ synthesis, and ORP5 has been shown to modulate levels of a PI(4,5)P_2_ reporter at the PM (Sohn, et al., 2018; Ghai, et al., 2017). Hence, the delay in BC formation upon ORP5 overexpression may be due to reductions in PI4P and PI(4,5)P_2_ at the PM. During normal development and maturation, ORP function at extensive ER-PM contacts may limit PI4P and PI(4,5)P_2_ levels in the adherens zona and specify a boundary for the apical domain (Figure 7b). Alternatively, ORP5 overexpression may restrict BC development and size by alterations in PS distribution or metabolism. Increased formation of ORP5-mediated ER-PM contacts could result in the inappropriate delivery of PS to basolateral domain and thus dilution of downstream PS effector proteins that are normally confined to the apical domain. In addition, ORP5 is proposed to function at ER-mitochondrial contacts (Galmes, et al., 2016) that are implicated in lipid metabolism including PS to PE conversion in hepatocytes (Vance, 1990). Similarly, ORP5 has been implicated in lipid droplet size control at ER-lipid droplet contacts (Du, et al., 2020) and thus ORP5 overexpression may alter ER-lipid droplet triacylglycerol lipid channelling and phospholipid (PS) synthesis in the ER. However, expression of a mutant ORP5 protein lacking its PH domain had no effect on BC development or apical aspect ratio (Figure 5 and Figure 5 – Figure supplement 1). The ORP5 PH domain is necessary for ORP5 localisation and function at ER-PM contacts (Sohn, et al., 2018; Chung, et al., 2015), but the ORP5 PH domain is not implicated in ORP5 targeting to ER-mitochondrial or ER-lipid droplet contacts (Du, et al., 2020; Galmes, et al., 2016). Taken together, these results suggest that ORP5 activity at ER-PM contacts defines apical domain size.

### Extensive ER-PM contacts are associated with lateral domains

While ORP5 restricted apical domain development and size, ORP5 in turn appeared to promote or stabilise cell-cell junctions (PM apposition stage between HepG2 couplets, Figure 5). As such, regulation of membrane lipid and Ca^2+^ dynamics at extensive ER-PM contacts may control junctional complexes in the lateral domains (adherens zona). For example, TJ formation and stability are proposed to be modulated by membrane lipid composition, particularly cholesterol content (Shigetomi, et al., 2018; Chen-Quay, et al., 2009; Nusrat, et al., 2000). ORP family members perform diverse biological functions including cholesterol transport (Olkkonen, 2015; Raychaudhuri and Prinz, 2010), suggesting potential roles of ORP-mediated ER-PM contacts in TJ positioning via cholesterol homeostasis. It will then be interesting to address roles of ORP family members and additional lipid transfer proteins in the formation and remodelling of other junctional complexes (adherens junctions and desmosomes) as well as the planar PM morphology that is observed at extensive ER-PM contacts within the lateral domain.

In conclusion, we have uncovered an unexpected role of the cortical ER network and ER-PM contacts in the regulation of epithelial cell polarity via phospholipid regulation. Two morphologically distinct ER-PM contacts are conserved in epithelial cells: rare punctate contacts at the apical domain and extensive contacts in the lateral domain. In addition, E- Syt1-mediated contacts may promote apical polarity at an early stage, whereas ORP5- mediated contacts may specify the apical boundary at later stages of cell development. Secretory vesicles also deliver phospholipids to the apical domain (Stoops and Caplan, 2014), and it will therefore be important to understand how non-vesicular transport at ER-PM contacts and polarised secretion are coordinated to specify PM domain identity. As such, the role of ER-PM contacts in cell polarity should not be considered independently from well-established pathways, but as an overlooked regulatory system that coordinates with known cell polarity networks. Beyond apico-basal polarity and epithelial cell development, it will also be exciting to examine roles of ER-PM contacts in vital hepatocyte functions including bile production and secretion, cholesterol homeostasis, steroid metabolism, insulin signalling, and glucose homeostasis. As our findings suggest that the arrangement of ER-PM contacts is conserved in epithelial cells, it will be interesting to elucidate physiological roles of ER-PM contacts in additional epithelial tissues as well. Finally, it is intriguing to consider whether remodelling of ER-PM contacts (for example, switching between E-Syt and ORP activities) is involved in epithelial cell and tissue remodelling during normal development and disease such as the epithelial to mesochymal transition during cancer progression.

## Materials and Methods

### Reagents

Plasmids and antibodies used in this study are listed in the Supplemental Tables.

### Acquisition of mouse liver samples

Mouse liver samples used for EM analysis were obtained from a previous study (Hanley, et al., 2017). Briefly, male mice were sacrificed at age 14 weeks and livers were harvested as described (Hanley, et al., 2017). For the Hanley et al. 2017 study, all animals were housed in accordance with the UK Home Office guidelines and the tissue harvesting was carried out in accordance with the UK Animal (Scientific Procedures) Act 1986. No additional animals were used during the course of the current study.

### Transmission electron microscopy (TEM) of hepatocytes from mouse liver and mIMCD spheroids

A piece of liver (approximately 8mm × 8mm × 3mm) was injection fixed with warm 1.5% glutaraldehyde in 1% sucrose 0.1M sodium cacodylate and left up to 20 minutes. Once firm and fixed, the tissue was cut into 100µm thick slices using a vibrating microtome. 2D hepatocyte and 3D IMCD cultures were fixed with 2% EM grade formaldehyde (TAAB) and 1.5% EM grade glutaraldehyde (TAAB) in either PBS or 0.1M sodium cacodylate. All samples were incubated for 1 hour in 1% osmium tetroxide/1.5% potassium ferricyanide at 4°C before being treated with 1% tannic acid in 0.05M sodium cacodylate (45 mins). Samples were then dehydrated through an ethanol series and embedded in Epon resin (TAAB). Ultrathin sections, 70nm, were collected on formvar coated slot grids using an Ultramicrotome (Leica), stained with lead citrate before being imaged using an 120kV Transmission electron microscope (Thermo Fisher Scientific) and images were captured using iTEM software and a Morada CCD camera (OSIS).

### Image analysis

#### Quantification of ER-PM contacts in tissues and organoids

EM images were formatted and analysed using Fiji (Schindelin, et al., 2012) and its inbuilt Straighten plugin (Kocsis, et al., 1991). See Figure 1 – Figure supplement 1 for the analysis workflow that quantified the PM category and physical parameters of the ER-PM contacts in hepatocytes and IMCD spheroids. Distances of 20 nm or less between the PM and ER were scored as contacts.

#### Segmentation of ultrastructure

Serial micrographs were stacked and aligned using the ImageJ StackReg Registration plugin (Thevenaz, et al., 1998) and manual segmentation was done using Amira (Thermofisher Scientific). The voxels of PM, ER, and ER-PM contacts were manually traced based on their electron density, morphological features, and the continuum of associated physical parameters.

### Cell culture and maintenance

HepG2 cells were cultured and maintained in Dulbecco’s Modified Eagle Medium/Nutrient Mixture F-12 (DMEM/F12; Life Technologies), supplemented with 10% (v/v) fetal bovine serum (FBS, Life Technologies), 100 U/mL of penicillin and 100 μg/mL of streptomycin at 37°C and 5% CO_2_. The cells were grown in a humidified incubator and passaged in a 1:15 ratio every 3 days or upon 90% confluency.

mIMCD cells were cultured following the protocol described in (Banushi, et al., 2016). For IMCD spheroid formation, the cells were cultured on an 8-well chamber slide (Lab-Tek) in DMEM/F12 supplemented with 2% FBS. The cells were sandwiched with Geltrex according to the manufacturer’s protocol (Life Technologies), and 2 × 10^4^ cells were seeded per well and grown at 37°C for 4 days to form spheroids.

### DNA Transfection

HepG2 cells were seeded at a density of 65,000 - 75,000 cells per 35mm dish (day 0) and grown overnight before a 24h transfection. Plasmids were introduced using Lipofectamine 3000 (Life Technologies) according to the manufacturer’s instructions. 0.5 μg of DNA plasmids expressing various lipid biosensors and/or proteins of interest was used per dish. Thereafter, confocal imaging or cell lysis was subsequently done (day 2).

### Confocal microscopy

Fluorescent microscopy was performed on a TCS SP5 microscope with Leica Application Suite: Advanced Fluorescence (LAS AF) software. For live imaging, cells were kept at 37°C with 5% CO2 in the Pe-Con stage-top incubation system throughout the imaging process and a 63x oil-immersion objective (NA=1.4) was used for image acquisition. Alternatively, the image was acquired using an Ultraview Vox spinning disc microscope operating on a Volocity interface with a C9100-13 EMCCD camera as the detector. A 60x oil-immersion objective (NA=1.4) was used.

### Immunofluorescence (IF) microscopy

For immunofluorescence microscopy, cells were fixed with 4% paraformaldehyde solution for 20 minutes, permeabilised using 0.5% saponin/3% bovine serum albumin (BSA) and then probed with desired primary and fluorescent-conjugated secondary antibodies. The cells were washed with PBS between each incubation step. Primary antibodies were incubated at 4°C overnight and secondary antibodies were incubated at room temperature for an hour, with the dilutions following manufacturer’s recommendation. The probed cells were kept in 2.5% (w/v) DABCO in PBS. When required, the cells were stained with 1 µg/ml DAPI to visualise the nuclei.

### SDS-PAGE and Immunoblotting

Cells were harvested using a cell scraper and were lysed with 200 μl/well of 0.1% CHAPS lysis buffer (50 mM Tris-HCl, 150 mM NaCl and 1 mM EDTA) supplemented with a Mini Protease inhibitor cocktail (Roche). The cell lysate was incubated on ice for 15 mins with occasional mixing before centrifugation at 20,000 g at 4°C for 10 min. The supernatant was collected and denatured in an SDS sample buffer at 99°C for 10 min before resolving on an 8% or 10% SDS-PAGE gel. Resolved proteins were then transferred to a supported nitrocellulose transfer membrane (GVS Filter Technology) in 10% methanol transfer buffer (Geneflow). The membrane was blocked against nonspecific antibody binding using 5% milk in Tris-buffered saline (TBS) containing 0.05% Tween 20 (TBST) for at least 1 hour and then probed with primary antibodies at 4°C overnight and horseradish peroxidase (HRP)- conjugated secondary antibodies at room temperature for 1 hour. Chemiluminescence was visualised using the ImageQuant LAS 4000 system and exported as digitalised images.

## Acknowledgements

We thank Tamas Balla, Pietro De Camilli, Greg Fairn, Jennifer Liou, and Rob Yang for reagents. We are grateful to Nathaniel Soh, Joanna Hanley, Rebecca Fiadeiro, and Ania Straatman-Iwanowska for expert technical assistance. We also thank Emily Eden, Tim Levine, and members of the Stefan lab for helpful discussions. CJS is supported by MRC funding to the MRC LMCB University Unit at UCL, award code MC_UU_00012/6. JJB is supported by MRC funding to the MRC Laboratory of Molecular Cell Biology at UCL, award code MC_U12266B. PG is supported by funding from the European Research Council, grant code ERC-2013-StG-337057.

## Author Contributions

GHCC, JJB, ML, PG, and CJS designed the study and experiments; GHCC and JJB performed experiments; GHCC, JJB, and CJS analysed the data; GHCC and CJS wrote the manuscript. All authors discussed the results and commented on the manuscript.

---

**Supplemental Movie 1**.

The PM, ER, and ER-PM contacts in the apical domain and lateral H/H contact domains (adherens zona) were segmented in EM serial sections. The subsequent 3D reconstruction displays the PM (cyan), ER (yellow) and ER-PM contacts (magenta).

**Supplemental Movie 2**.

Segmentation of the PM (cyan), ER (yellow), and ER-PM contacts (magenta) in serial EM sections of the basal domain and subsequent 3D reconstruction.

**Supplemental Movie 3**.

Serial TEM sections of a developing bile canaliculus between a HepG2 couplet. Note the abundance of microvilli in the apical domain and punctate ER-PM contacts at the base of microvilli (an example can be clearly seen in the last frame of the movie).

## Supplemental Tables

**Table S1.**
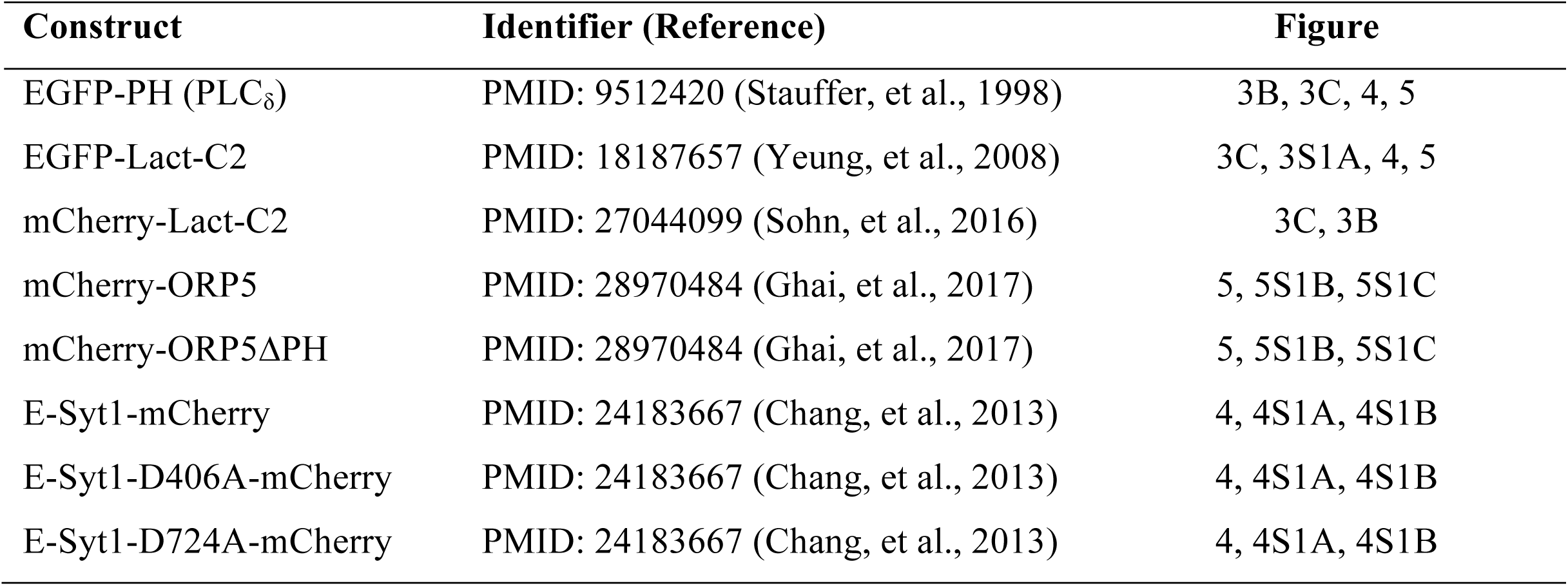
Plasmids used in this study

| Construct | Identifier (Reference) | Figure |
| --- | --- | --- |
| EGFP-PH (PLCδ) | PMID: 9512420 (Stauffer, et al., 1998) | 3B, 3C, 4, 5 |
| EGFP-Lact-C2 | PMID: 18187657 (Yeung, et al., 2008) | 3C, 3S1A, 4, 5 |
| mCherry-Lact-C2 | PMID: 27044099 (Sohn, et al., 2016) | 3C, 3B |
| mCherry-ORP5 | PMID: 28970484 (Ghai, et al., 2017) | 5, 5S1B, 5S1C |
| mCherry-ORP5ΔPH | PMID: 28970484 (Ghai, et al., 2017) | 5, 5S1B, 5S1C |
| E-Syt1-mCherry | PMID: 24183667 (Chang, et al., 2013) | 4, 4S1A, 4S1B |
| E-Syt1-D406A-mCherry | PMID: 24183667 (Chang, et al., 2013) | 4, 4S1A, 4S1B |
| E-Syt1-D724A-mCherry | PMID: 24183667 (Chang, et al., 2013) | 4, 4S1A, 4S1B |

**Table S2.** Primary antibodies used in this study

| Antibody | Identifier (Source) | Figure |
| --- | --- | --- |
| Anti-OSBPL5 | ab59016 (Abcam) | 5S1B |
| Anti-CEA | a1105 (DAKO) | 3B |
| Anti-MRP2 | ab3373 (Abcam) | 3B |
| Anti-Par3 | 07-330 (Millipore) | 3S1A |
| Anti-mCherry | ab125096 (Abcam) | 4S1A |
| Anti-GAPDH, HRP | PA1-987-HRP (Invitrogen) | 4S1A |

## Figure and Figure Supplement Legends

**Figure 1 - Figure supplement 1.**
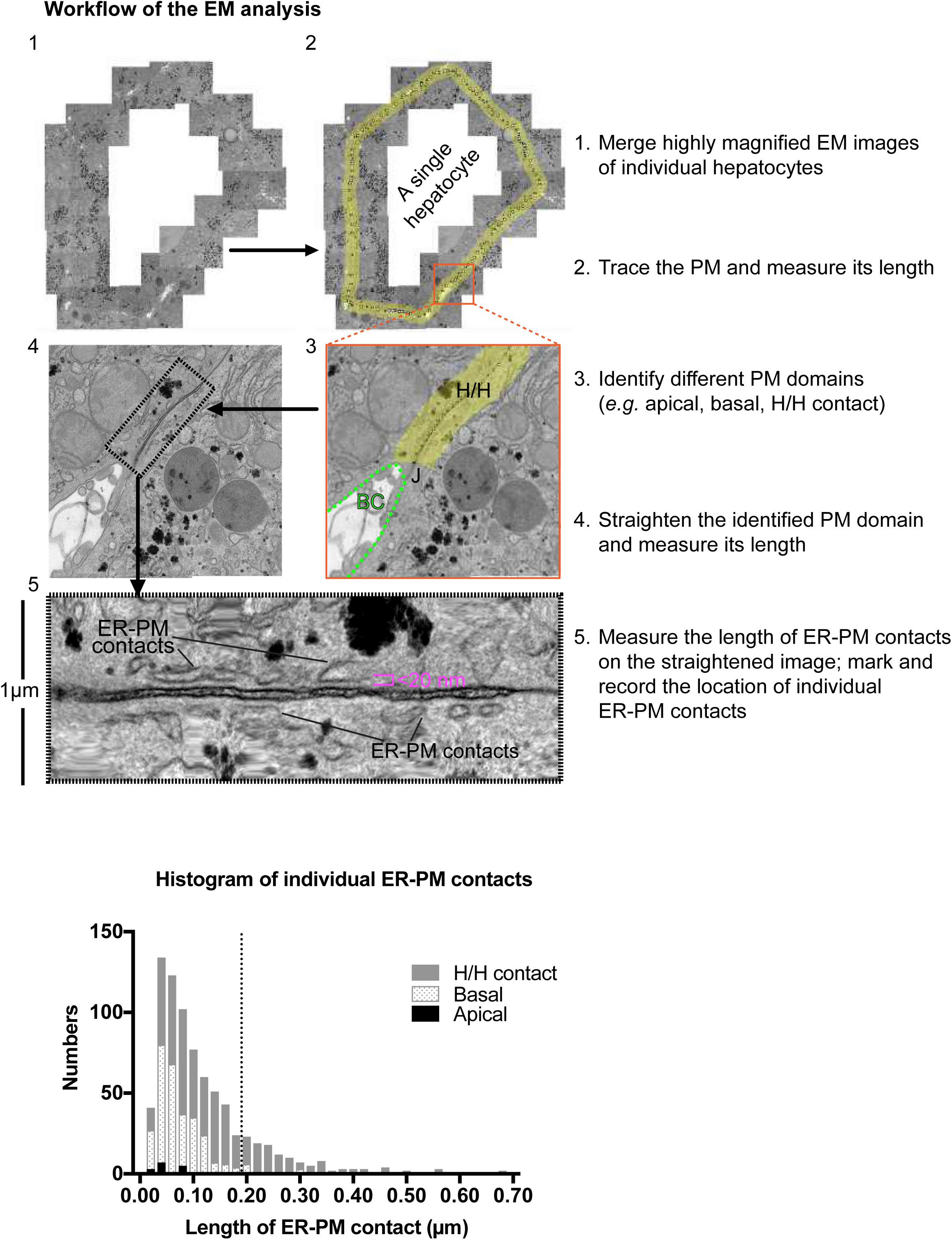
**Top panel:** The image analysis workflow used to quantify the PM category and the length and distribution of ER-PM contacts in hepatocytes. The PM and associated cortical region of the hepatocyte (yellow) were straightened using the Fiji straighten plugin (Kocsis, et al., 1991) so that it allows assessment of the distance between the PM and the ER. Any tethering of the ER to the PM ≤ 20 nm was classified as ER-PM contact, in accord with criterion employed in a previous analysis (Hsieh, et al., 2017). **Bottom panel:** The histogram of the length distribution of all individual ER-PM contacts in the 9 hepatocytes analysed from 3 animals.

**Figure 1 - Figure supplement 2.**
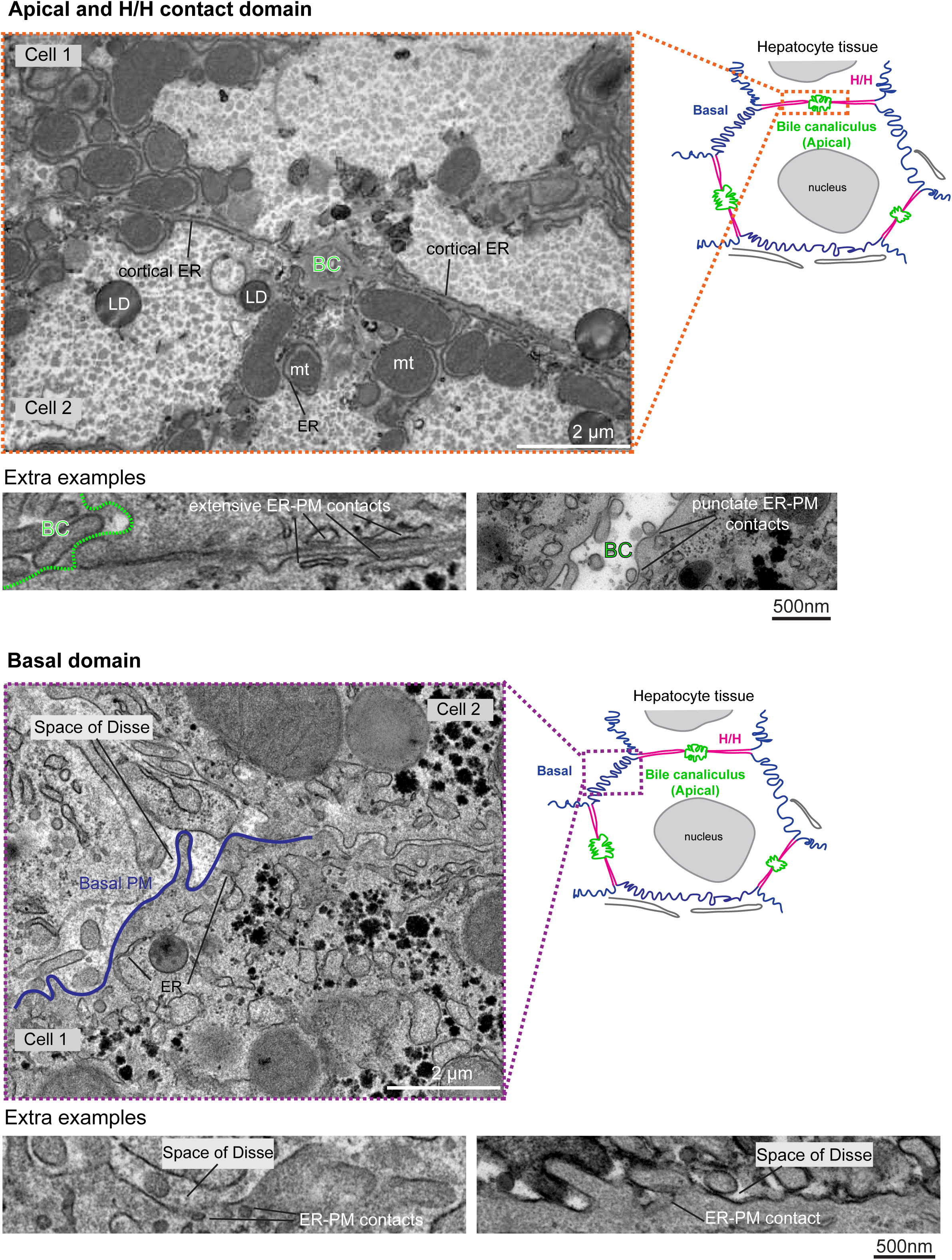
Representative electron micrographs of different PM domains in hepatocytes. Extra examples highlight the differences in length and morphology of the ER-PM contacts in the apical and H/H contact PM domains.

**Figure 1 - Figure supplement 3.**
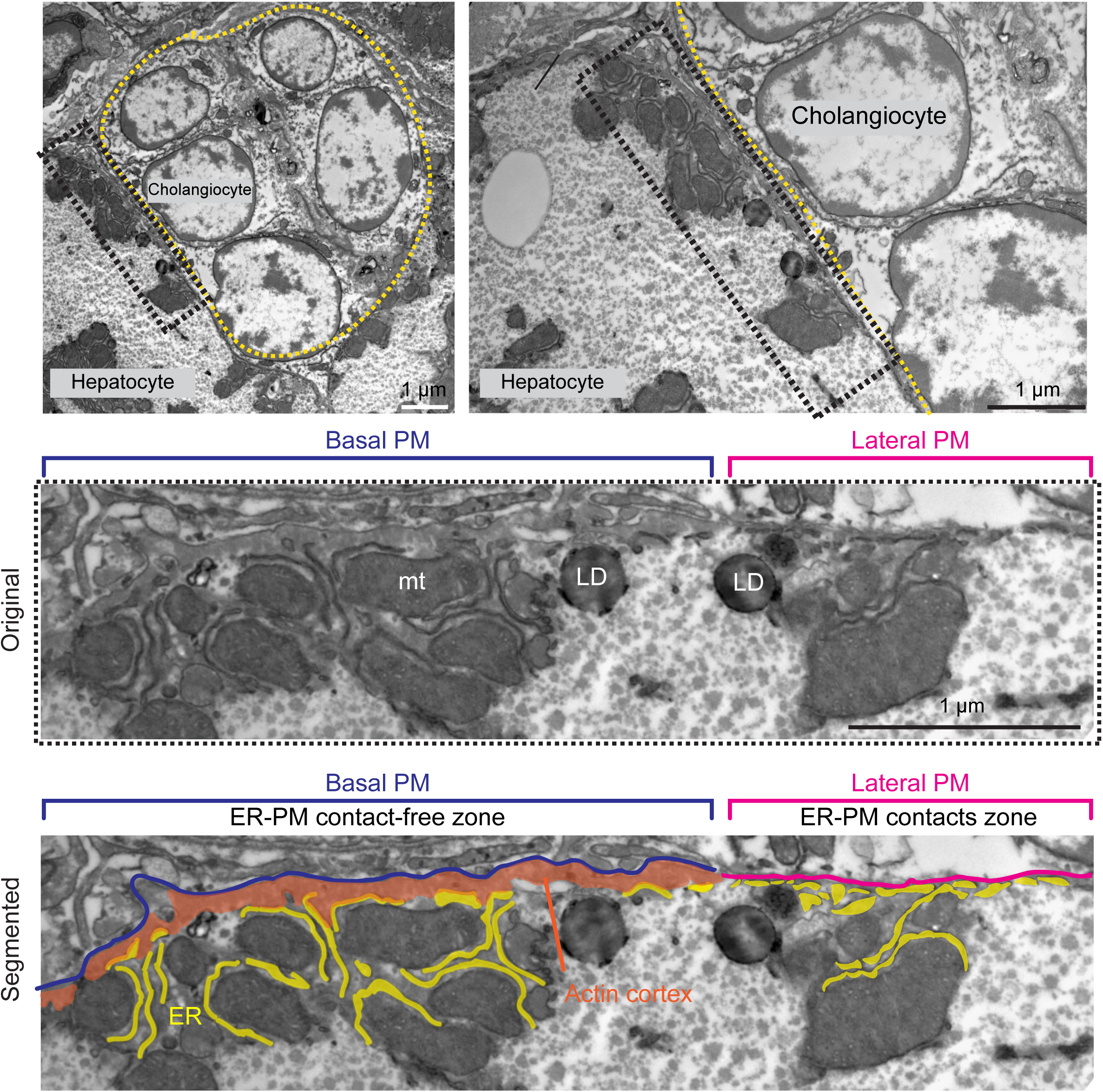
**Top and middle panels:** Electron micrographs of the interface between a hepatocyte and cholangiocytes. The yellow dashed line marks the boundary of the cholangiocyte bundle and the black box locates the ROI in the hepatocyte juxtaposed to the cholangiocytes. The basal and lateral PM are coloured in blue and magenta respectively. **mt:** mitochondria; **LD** lipid droplets. **Bottom panel:** A segmented micrograph depicts the organisation of the PM, ER and actin in the hepatocyte-cholangiocyte interface. Note the presence of the actin cortex and the absence of ER-PM contacts in the basal PM.

**Figure 2 - Figure supplement 1.**
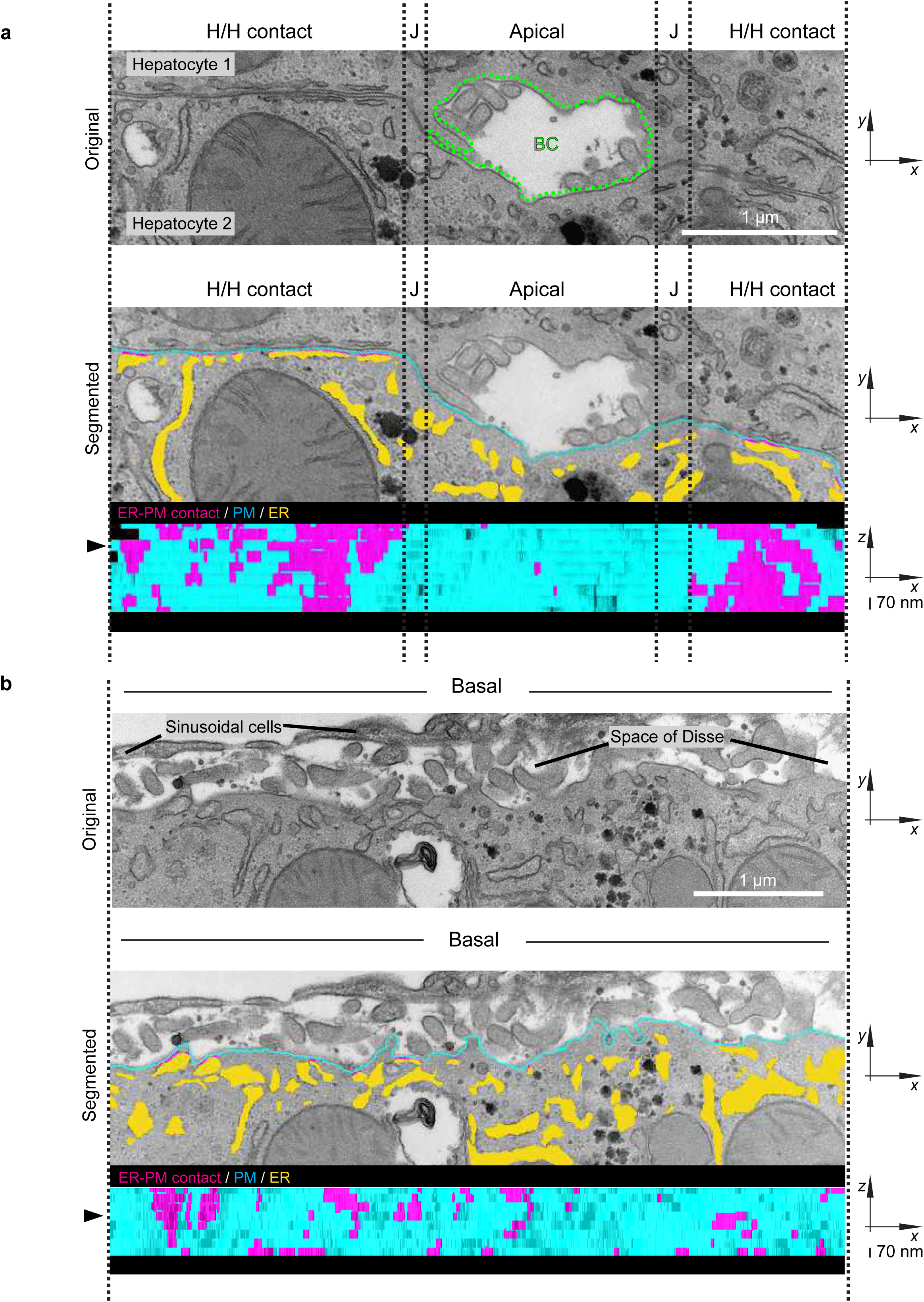
Additional examples of the segmentation analyses and 3D model projections of ER-PM contacts in different PM domains in hepatocytes; see the main figure for an identical figure description.

**Figure 2 - Figure supplement 2.**
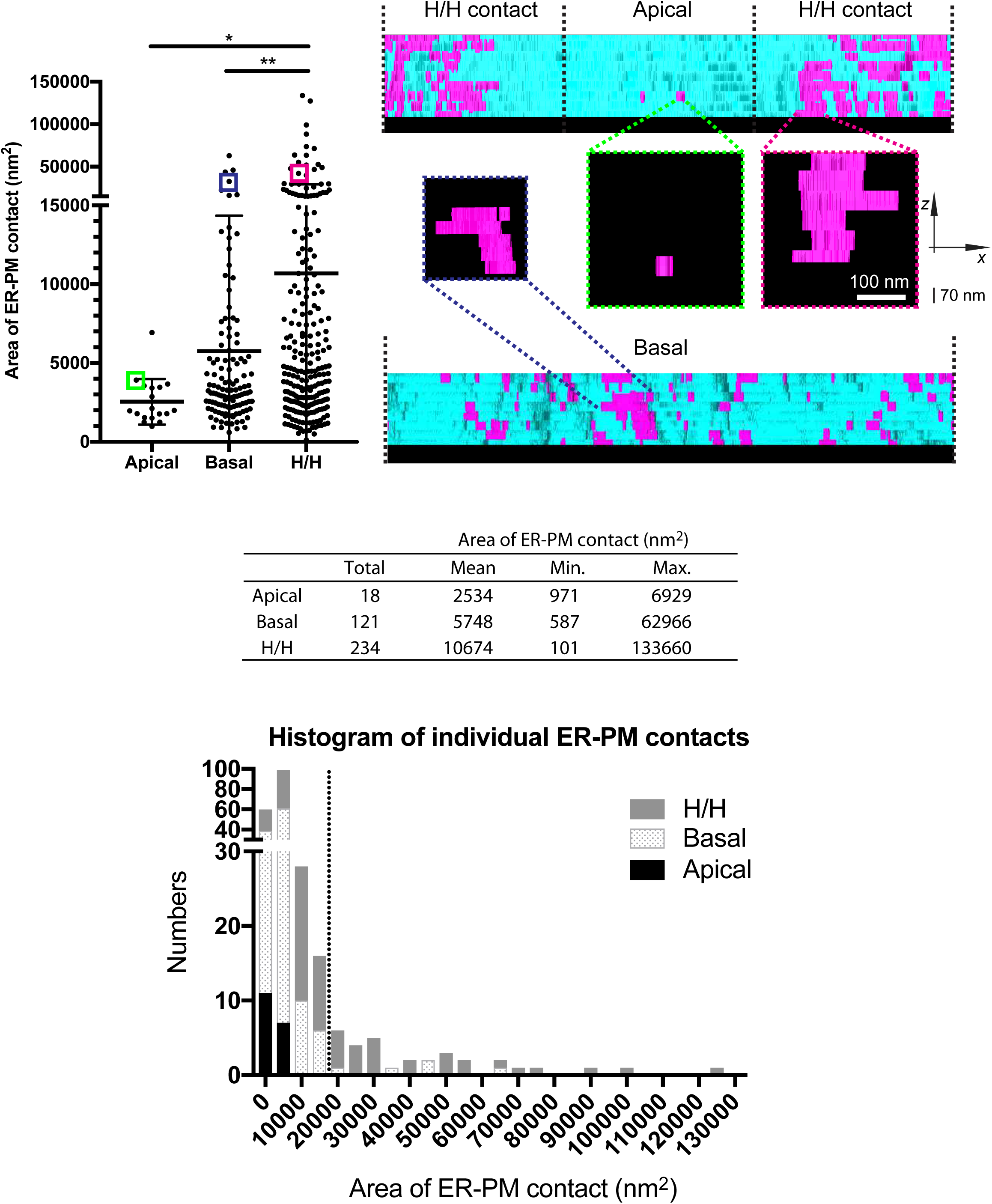
Quantification of the segmentation analyses. The scatter dot plot shows the area of the ER-PM contacts (mean ± SD) derived from the *x-*z projections of the reconstructed 3D models (N=3 apical and H/H contact domain; N=2 basal domain; approximately equal numbers of voxels per PM domain were compared). Data were analysed by one-way ANOVA followed by a Tukey’s multiple comparison test: **p<0.01; *p<0.1. Panels on the right serve as a visual comparison of the largest ER-PM contact in each PM domain from Figure 2. The area of the selected ER-PM contacts is indicated by the coloured boxes in the plot. The table summarises the quantity and range of the area of the ER-PM contacts, and the histogram reveals their area distribution in each PM domain.

**Figure 3 - Figure supplement 1.**
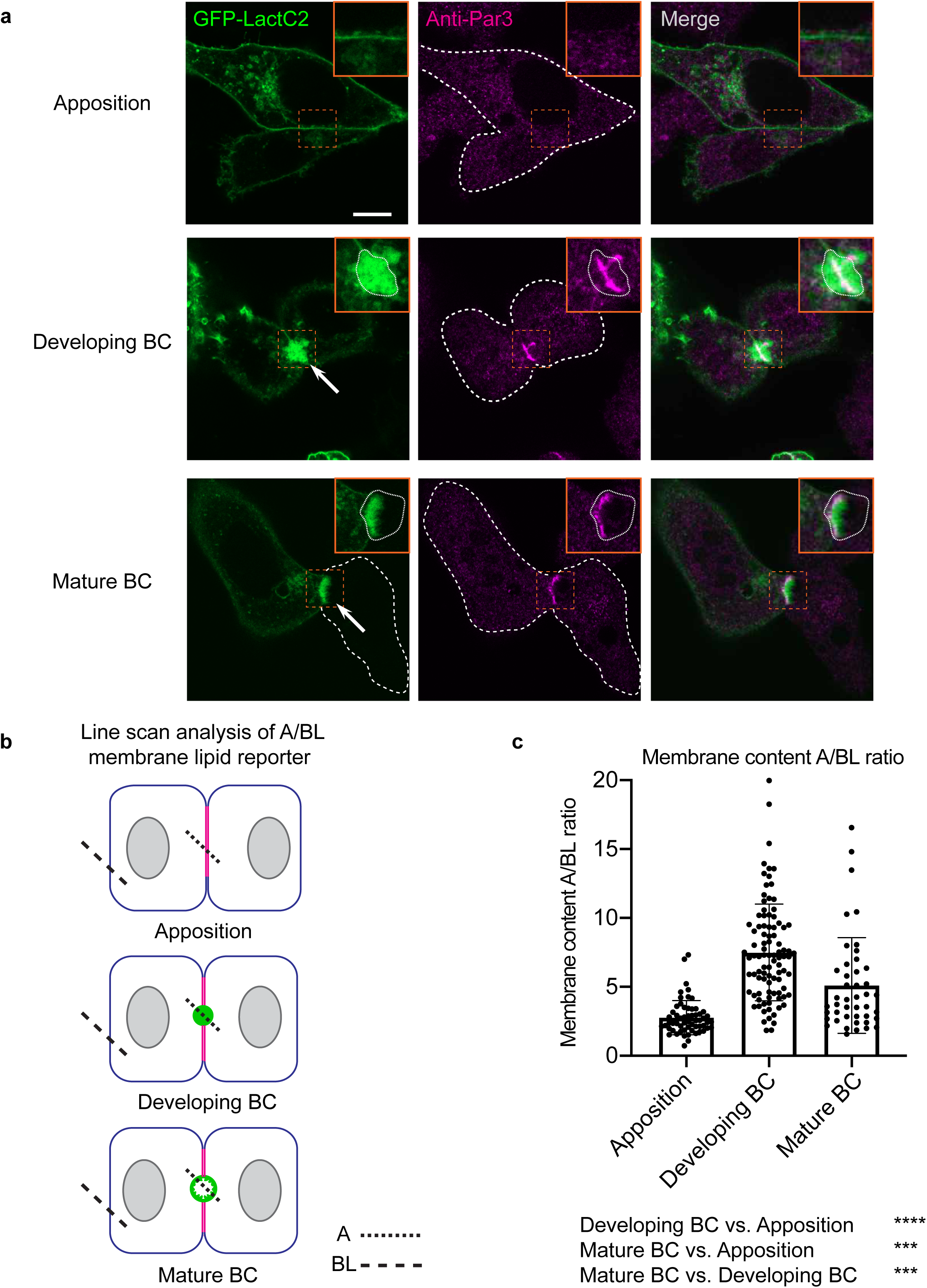
**(a)** Confocal images (mid-sections) of HepG2 cells overexpressing the GFP-Lact-C2 PS reporter and stained with the anti-Par3 polarisation marker. The arrows indicate the position of the developing or matured BC, of which the boundaries are outlined in the 2X magnified inlets. Scale bar = 10 *μ*m. **(b)** A diagram illustrates the line scan approach we used to measure the apical to basolateral lipid intensity of HepG2 couplets. For HepG2 couplets in the apposition stage the intensity of the mid-point at the cell-cell contact was taken as the apical value, since this is often the site where developing BC will form. **(c)** A scatter dot plot shows the apical to basolateral intensity ratio of the CellMask PM dye during BC development. These values were used to normalise levels of the PS and PI(4,5)P_2_ reporters relative to the amount of total membrane in the apical domain (see Figure 3c). Data are shown as mean ± SD and were analysed by one-way ANOVA (Tukey’s multiple comparison test: ****p<0.0001; ***p<0.001; N=4 independent experiments).

**Figure 4 - Figure supplement 1.**
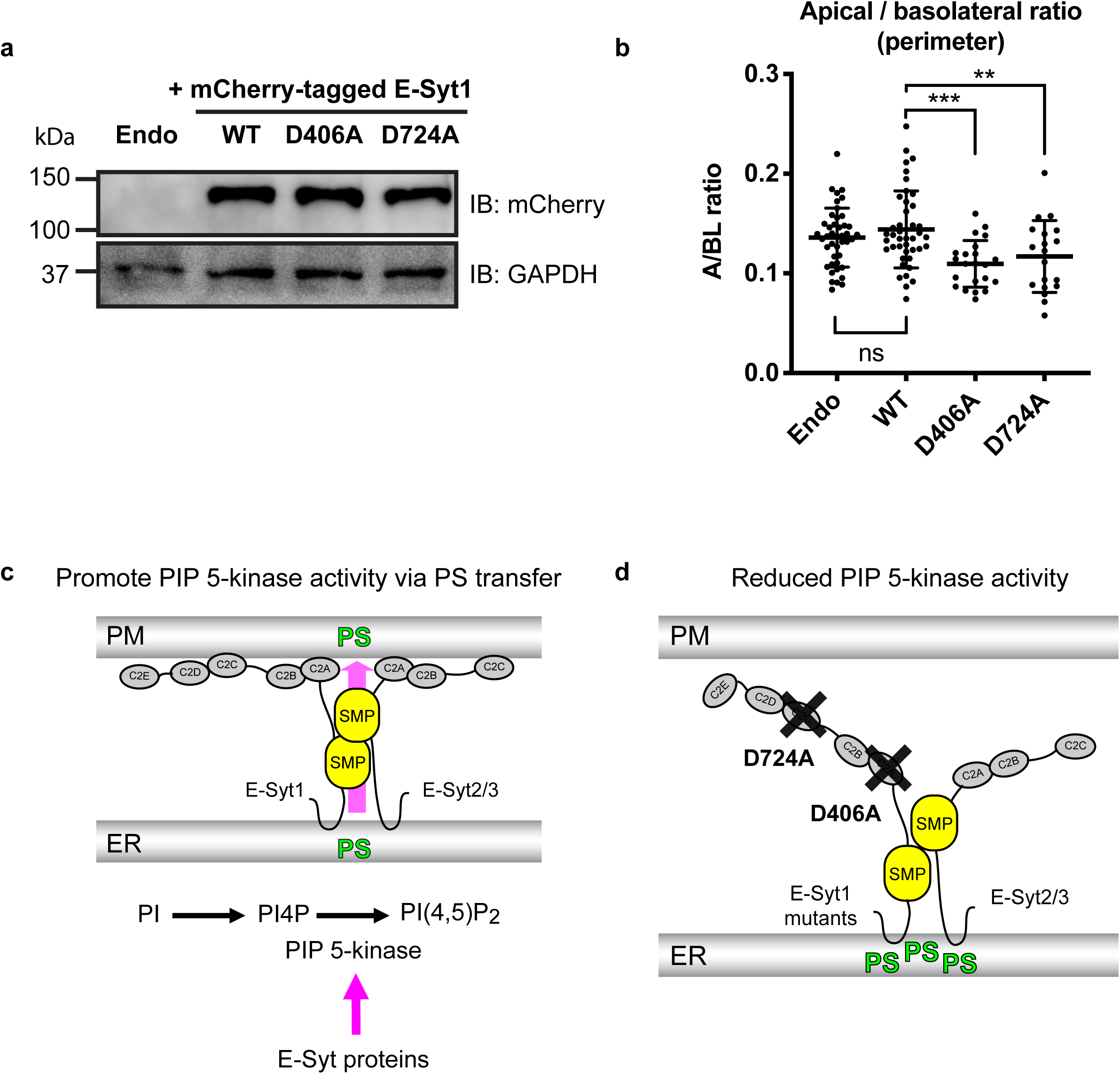
**(a)** An immunoblot reveals the protein expression levels of the mCherry-tagged E-Syt1 variants in HepG2 cells. **(b)** The apical to basolateral ratio of PM perimeter in mature HepG2 couplets expressing different E-Syt1 variants and a lipid reporter. Data are shown as mean ± SD and were analysed by one-way ANOVA (Dunnett’s test: ***p<0.001; **p<0.01 N=6 independent experiments). **(c)** A speculative role of E-Syt proteins in promoting PIP 5-kinase activity, thus PI(4,5)P_2_ synthesis, by PS transport via their dimerised SMP domains. Whether putative PS transport may occur at ER-PM contacts by a ‘shuttle’ or a ‘tunnel’ mechanism is not addressed in this study. See the main text for alternative explanations for the role of E- Syt1 in PS and PI(4,5)P_2_ regulation at the apical membrane. **(d)** A diagram suggesting how overexpression of the D406A or D724A E-Syt1 mutants may form less stable ER-PM contacts with the endogenous E-Syt1/2/3 proteins, resulting in impaired PS delivery to the PM and accumulation of PS at intracellular membrane compartments.

**Figure 5 - Figure supplement 1.**
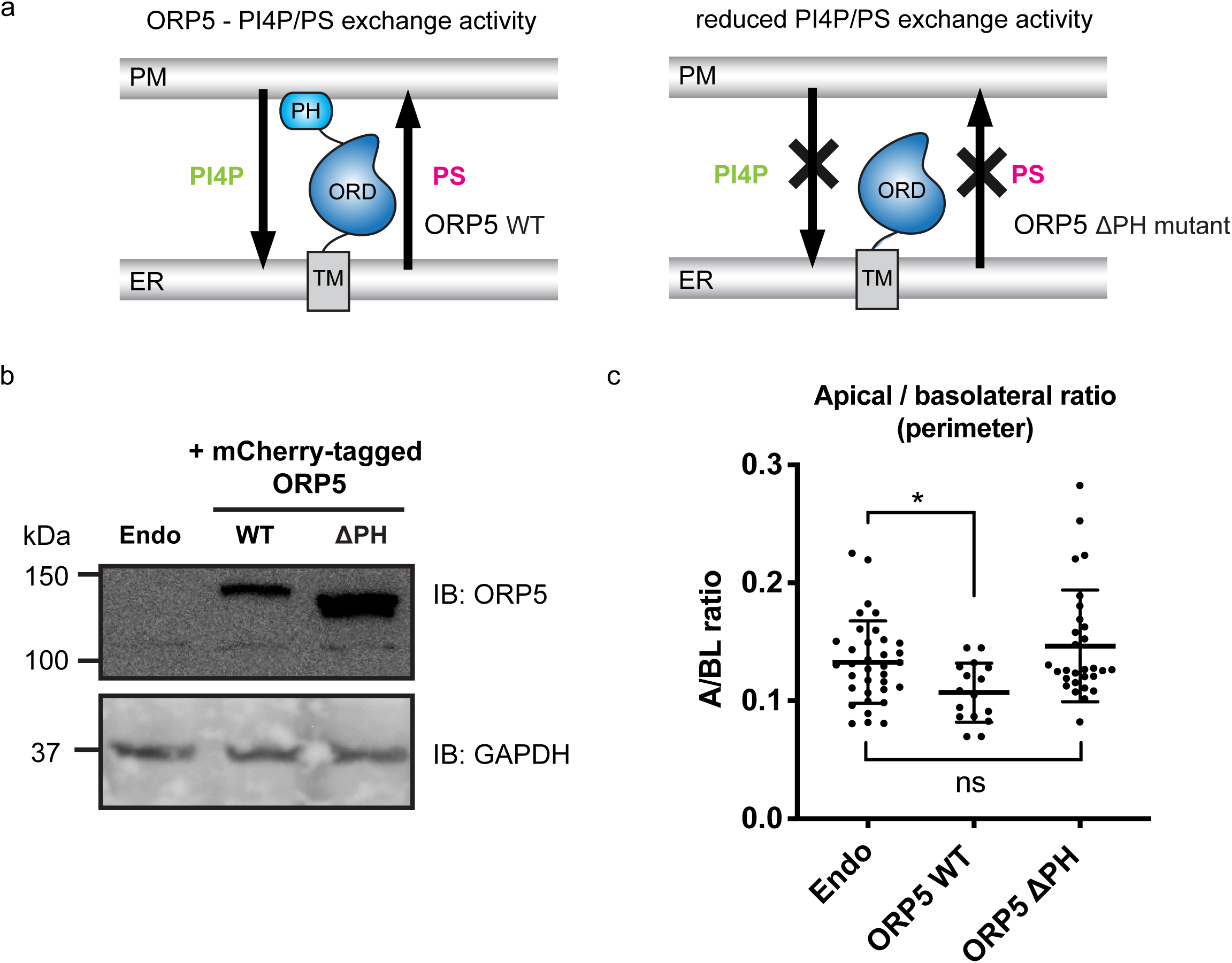
**(a)** The ORP5 protein is proposed to carry out PI4P/PS exchange at ER-PM contacts, and deletion of the ORP5 PH domain impairs ORP5 localisation and lipid exchange at ER-PM contacts. **(b)** An immunoblot reveals the protein expression levels of the mCherry-tagged ORP5 and ΔPH mutant in HepG2 cells. **(c)** The apical to basolateral ratio of PM perimeter in matured HepG2 couplets expressing the ORP5 variants and a lipid reporter. Data are shown as mean ± SD and were analysed by one-way ANOVA (Dunnett’s test: *p<0.1 N=3 independent experiments).

**Figure 6 - Figure supplement 1.**
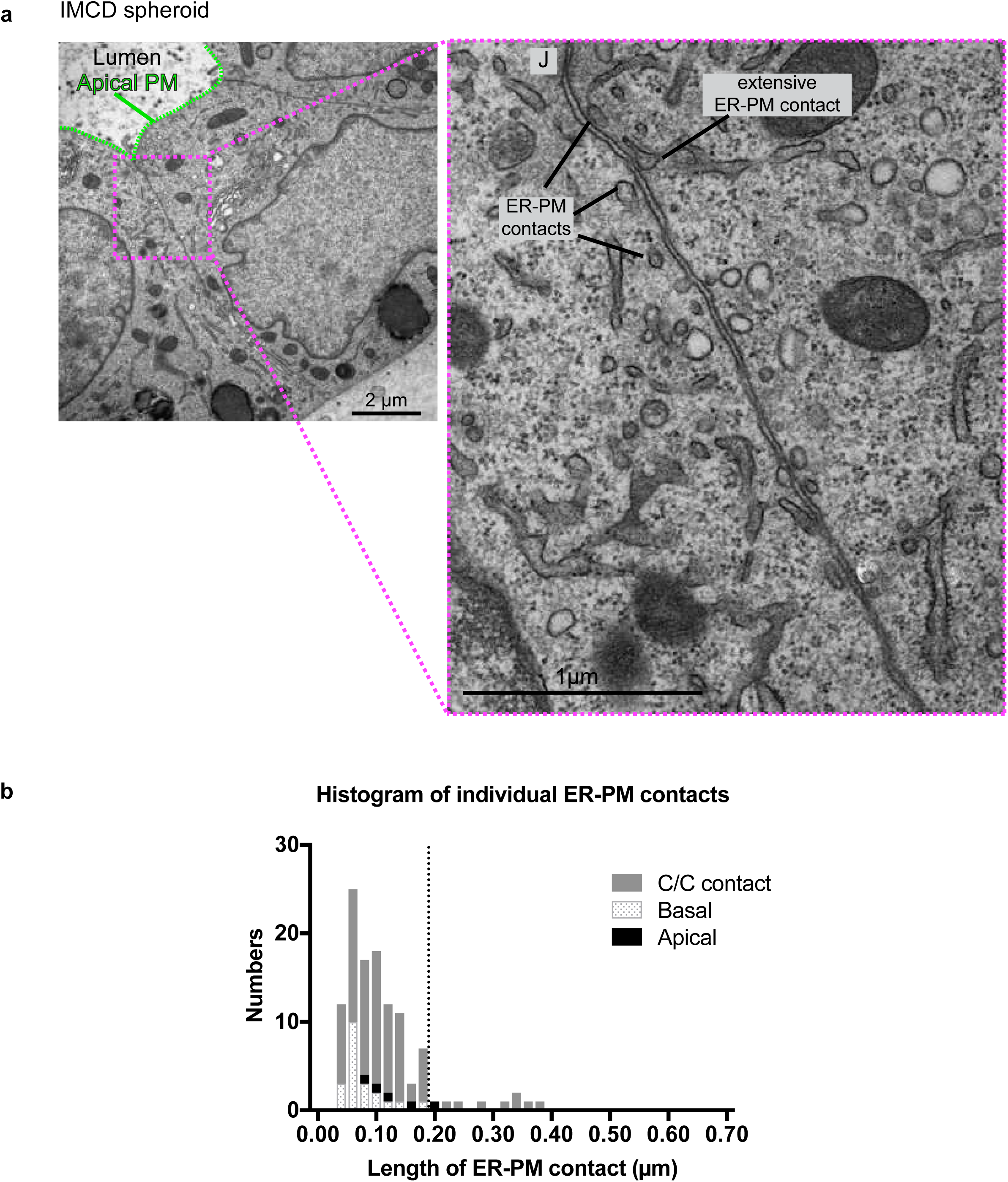
**(a)** Electron micrographs showing the ultrastructure of two apposed IMCD cells within a spheroid. Note the presence of multiple ER-PM contacts along the lateral PM, and also the presence of an extensive ER-PM contact in the vicinity of the cell junction. **(b)** A histogram reveals the length distribution of ER-PM contacts in the IMCD spheroids we analysed.

